# Temperature-Dependent Rotamer Population Shifts Govern Tryptophan Fluorescence in Proteins

**DOI:** 10.64898/2026.05.22.726722

**Authors:** I-Shan Hsu, Yun-Chu Chou, Wei-Hsiang Wang, Min-Yeh Tsai

**Affiliations:** Department of Chemistry and Biochemistry, National Chung Cheng University, Minhsiung, Chiayi 621301, Taiwan; Center for Nano Bio-Detection, National Chung Cheng University, Minhsiung, Chiayi 621301, Taiwan; Department of Chemistry, Tamkang University, New Taipei City 251301, Taiwan

## Abstract

Intrinsic tryptophan fluorescence is widely used as a sensitive reporter of protein conformational dynamics, yet the molecular origin of its temperature-dependent modulation remains unclear. Here we investigate the conformational dynamics of Trp134 in bovine serum albumin (BSA) using molecular dynamics (MD) simulations, free-energy calculations based on umbrella sampling and WHAM, quantum mechanical (QM) calculations, and QM/MM approaches. MD simulations show that the global structure of BSA remains stable while temperature induces a gradual population shift from the Ia^+^ to the Ia^−^ rotamer. The corresponding free-energy landscapes reveal that this shift arises from subtle changes in basin stability and transition barriers along the rotameric coordinate. In contrast, standalone QM calculations on isolated tryptophan predict different energetic trends, highlighting the sensitivity of rotamer stability to electronic-structure treatments and environmental effects. QM/MM calculations partially reconcile these differences by incorporating the protein environment. Together, these results suggest that temperature reshapes the rotamer free-energy landscape of Trp134, leading to population shifts that modulate intrinsic tryptophan fluorescence in proteins.

## INTRODUCTION

Tryptophan fluorescence is a valuable tool for studying protein structure and function, offering insights into local environments and conformational changes.[1] The fluorescence characteristics of tryptophan are susceptible to various micro-environmental factors, including temperature, pH, and polarity.[2] This sensitivity has made tryptophan fluorescence widely used for studying protein folding and biomolecular interactions.[3] Despite its broad application, the molecular mechanisms behind tryptophan’s fluorescence behavior, especially the temperature dependence of fluorescence intensity, remain poorly understood.[4] Gaining a deeper understanding of these mechanisms is crucial for improving the use of tryptophan fluorescence as a reliable probe in molecular dynamics studies.

Tryptophan, an aromatic amino acid, contributes significantly to the intrinsic fluorescence of proteins. The indole ring of tryptophan absorbs light, undergoes excited-state relaxation, and emits fluorescence sensitive to the local environment and overall protein structure. Tryptophan residues exhibit distinct fluorescence lifetimes, typically categorized into short (*τ*_1_) and long (*τ*_2_) decay times, which are influenced by the local microenvironment and the conformational state of tryptophan.[5,6] These lifetimes can also be affected by environmental factors, including temperature, pH, and protein dynamics. Studies by Kaczor et al. have shown that the fluorescence properties of tryptophan in proteins are governed by its conformational isomers, or rotamers, specifically defined by the two key dihedral angles *χ*_1_ (C_COOH_–C_α_–C_β_–C_γ_) and *χ*_2_ (C_α_–C_β_–C_γ_=CD1), whose orientations (a/b/c and +/– conformations) are significantly influenced by intramolecular hydrogen bonds and electrostatic interactions involving the pyrrole and phenyl rings (See Fig.1).[7] These rotamers interconvert sensitively in response to changes in temperature, solvent conditions, and surrounding protein dynamics.[7] Consistent with this picture, time-resolved fluorescence studies of serum albumin have provided experimental evidence that discrete rotamer conformations contribute to fluorescence lifetime heterogeneity, supporting a direct connection between rotamer populations and fluorescence observables.[8]

**Figure 1.**
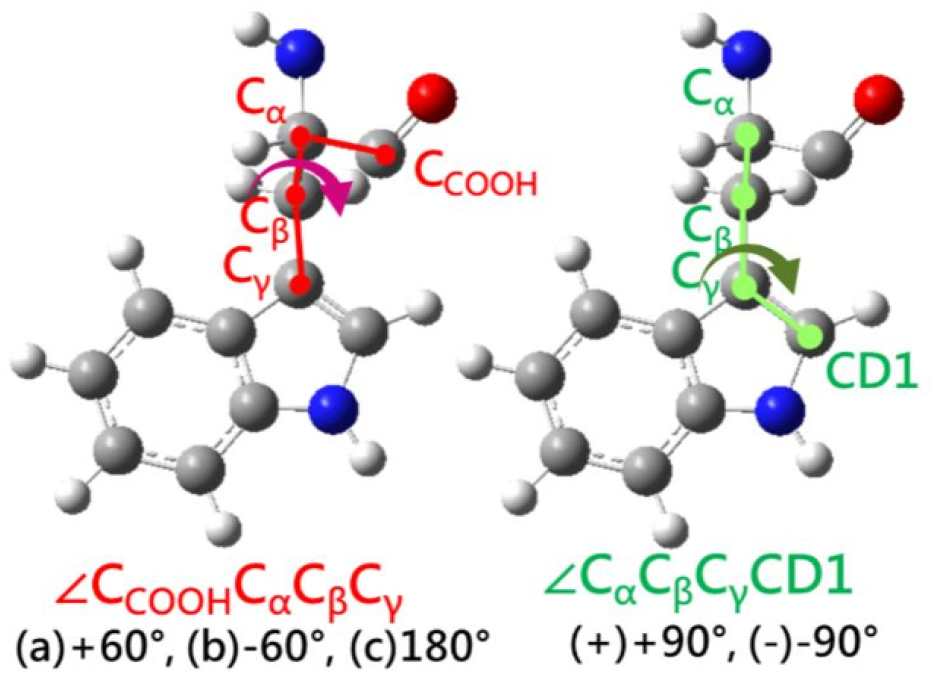
Definition of the two dihedral angle sets used to classify tryptophan rotamers in BSA. The first set (φ1 = ∠*C*_*COOH*_*C*_α_*C*_β_*C*_γ_) distinguishes three orientations: (a) +60°, (b) –60°, and (c) +180°. The second set (φ2 = ∠*C*_α_*C*_β_*C*_γ_*CD*1) distinguishes two orientations: (+) +90° and (–) –90°. Combinations of φ1 and φ2 yield six possible rotamers: Ia^+^ (60°, 90°), Ia^−^ (60°, −90°), Ib^+^ (−60°, 90°), Ib^−^ (−60°, −90°), Ic^+^ (180°, 90°), and Ic^−^ (180°, −90°).

One of the key observations in the study of tryptophan fluorescence is the temperature dependence of its emission intensity. For example, in the case of bovine serum albumin (BSA), the fluorescence of tryptophan residues such as Trp134 and Trp213 exhibits distinct temperature-sensitive behavior.[9,10] The fluorescence intensity of Trp134 decreases as the temperature rises, which has been attributed to temperature-induced conformational changes in the protein.[9] However, the exact molecular mechanism that underlies this phenomenon remains a subject of debate. Currently, several factors contribute to the temperature dependence. Photoionization is a major nonradiative pathway that competes with fluorescence in aqueous tryptophan, and its efficiency strongly depends on temperature.[11] Studies have shown that the efficiency of photoionization increases significantly as temperature rises. This is because the rate of photoionization accelerates, outcompeting radiative decay and thereby reducing fluorescence yield and lifetime. At a neutral pH (pH 7), an additional temperature-dependent intramolecular quenching process, likely involving side-chain conformational rearrangements and proton transfers, further contributes to fluorescence suppression.[11] In brief, these temperature-sensitive nonradiative processes explain the pronounced decrease in tryptophan fluorescence observed at elevated temperatures.

Alternatively, in a recent study, Chou et al. proposed a rotamers-based model[12] and demonstrated that tryptophan residues undergo thermal transitions that affect their fluorescence properties. The temperature dependence of tryptophan fluorescence is thought to arise from population shifts between different rotamers, with certain major rotamers corresponding to distinct fluorescence lifetimes, and these shifts are driven by changes in the relative stability of the rotamers at different temperatures.[12] Such behavior can be interpreted within the framework of population redistribution on a rugged free-energy landscape, where subtle energetic changes shift the equilibrium among metastable conformational substates and thereby modulate observable physical properties. [13] Previous studies have examined tryptophan rotamers using either quantum chemical calculations on isolated systems or molecular simulations within proteins.[7,14,15] However, how MD-, QM-, and QM/MM-based descriptions of rotamer stability compare—and whether they lead to consistent interpretations of temperature-dependent fluorescence—remains largely unexplored.

In the current study, we employ a combined approach of molecular dynamics (MD) simulations and quantum mechanical (QM) calculations to investigate this phenomenon. First, we demonstrate that the temperature-dependent fluorescence observed in bovine serum albumin (BSA) predominantly arises from the local rotameric dynamics of the Trp134 residue rather than significant changes in overall protein structure. Our MD simulations clearly illustrate population shifts among Trp134 rotamers. Second, we analyze the free energy profile (potential of mean force, PMF) governing the interconversion between Trp rotamers in BSA. The resulting free energy landscape aligns well with the temperature-dependent fluorescence predicted by the rotamer model. Additionally, we explore solvation effects during rotamer interconversion, highlighting the solvent environment’s critical role in stabilizing transition states and modulating energy barriers. Third, we compare the insights obtained from MD and QM approaches and identify discrepancies in their description of temperature-dependent fluorescence. To resolve these discrepancies, we perform QM/MM hybrid calculations, integrating the strengths of both methods to accurately characterize the free energy trend and evaluate the general role of tryptophan rotameric dynamics in fluorescence emission responses to temperature. Ultimately, this work validates the effectiveness of MD simulations in exploring local side-chain conformational dynamics affecting protein fluorescence and clarifies the molecular mechanism behind fluorescence sensitivity to temperature.

## METHODS

### MD simulations

All-atom molecular dynamics (MD) simulations were carried out using NAMD 2.14[16] with CUDA acceleration on RTX 2080Ti GPUs to investigate the dynamic behavior of fluorescent amino acid residues within proteins. The CHARMM27 force field was employed, and the simulation setup was prepared using VMD 1.9.3.[17] The initial structure of bovine serum albumin (BSA) was obtained from PDB ID: 4F5S.[18] The protein was solvated in a water box, and 17 Na^+^ ions were added to neutralize the system, yielding an ion concentration of 60 mM. For comparison under physiological conditions, an additional set of simulations was performed at 150 mM ion concentration (see Fig. S1 of the Supporting Information). The final simulation box contained 54,240 atoms (protein: 9,253; ions: 17; water: 44,970, equivalent to 14,990 molecules) with dimensions of 90 Å × 72 Å × 93 Å. Periodic boundary conditions (PBC) were applied, and long-range electrostatics were calculated using the Particle Mesh Ewald (PME) method. Prior to production runs, the system was energy-minimized for 100 steps. MD simulations were conducted with a 2 fs timestep and rigidBonds enabled. Temperature was maintained using a Langevin thermostat (damping coefficient: 1 ps^−1^), and pressure was controlled with the Langevin piston method (1 atm). Each trajectory was simulated for 480 ns, with five independent replicas per temperature, resulting in a total sampling time of 2.4 μs. Trajectories were saved every 10 ps. Four temperature conditions were examined: 303 K, 313 K, 323 K, and 353 K.

To complement the BSA simulations, additional MD simulations were performed on an isolated Trp134 residue under both gas-phase (vacuum) and solvated conditions with explicit water. The solvated system comprised 1806 atoms, including one Trp residue and 590 water molecules, whereas the gas-phase system contained only 24 atoms corresponding to a single Trp residue. These comparative simulations allowed us to separate the intrinsic conformational dynamics of Trp134 from environmental influences, thereby clarifying the roles of solvation and the protein matrix in modulating the fluorescence-related behavior of tryptophan.

### Global and local structural analysis

First, the overall RMSD of the BSA protein structure was calculated to observe the structural changes in BSA at different temperatures (303K, 313K, 323K and 353K). The RMSD is a measure of the average distance between atoms of two structures and is calculated using the formula:

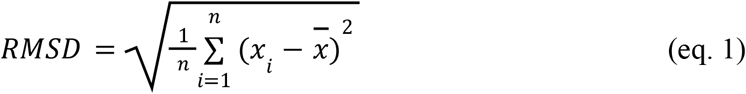

where *n* is the total number of protein backbone atoms, including N, O, C_α_ and C, *x*_*i*_ is the coordinate of atom *i* in the current structure, and 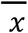 is the coordinate of atom *i* in the reference structure. The RMSD value gives the average deviation between the corresponding atoms of two proteins: the smaller the RMSD, the more similar the two structures. RMSD provides a measurement of the average structural fluctuation of a protein relative to a reference structure. It reflects how much the overall protein conformation deviates during the simulation, serving as a general indicator of structural stability under different conditions.

To investigate local structural dynamics, RMSD can also be applied to specific subdomains or secondary structure elements, as exemplified in many globular proteins.[19] In this study, a subdomain including Trp134 and three α-helices surrounding Trp134 (residues 15–31, 129–145, and 149–168) were selected. RMSD calculations were performed both on the entire subdomain and on individual α-helix to capture local conformational variations. Such localized RMSD analysis helps determine whether structural fluctuations occur preferentially in certain regions rather than the whole protein.

### The Rotamer model

Based on the studies of Sanz et al. and Kaczor et al., tryptophan rotamers have been classified into 17 types,[14] defined by the hydrogen-bonding patterns between the amino and carboxyl groups, as well as the backbone and side-chain dihedral angles.[7] In our case, since tryptophan is considered as a residue within proteins, only type I hydrogen bonding (N–H···O=C) is relevant. This restriction reduces the number of possible rotamers to six, each uniquely specified by a set of dihedral angles. A dihedral angle φ is defined by four atoms (A–B–C–D), where two planes are formed: one passing through atoms A, B, and C, and the other through atoms B, C, and D. The dihedral angle corresponds to the angle between these two planes. In proteins, dihedral angles describe both backbone conformations and side-chain orientations, making them essential parameters for characterizing conformational states such as rotamers.

To quantify the dihedral angles of the two tryptophan residues in BSA and analyze their rotameric flipping dynamics, we adopted two sets of dihedral angle definitions: ∠*C*_*COOH*_*C*_α_*C*_β_*C*_γ_, which can take values of approximately (a) +60°, (b) −60°, (c) +180°, and ∠*C*_α_*C*_β_*C*_γ_*CD*1, which adopts either (+) +90° or (−) −90°. These combinations define six rotamers observed in the experiments: Ia^+^ (60°, 90°), Ia^−^ (60°, −90°), Ib^+^ (−60°, 90°), Ib^−^ (−60°, −90°), Ic^+^ (180°, 90°), and Ic^−^ (180°, −90°). Fig.1 illustrates the definitions of these two dihedral angle sets used to characterize the rotamers.

Building on the dihedral definitions described above, we previously proposed a rotamer model linking the quantum yield (*I*_Φ_) to the free energy difference (Δ*F*) between two rotameric states, as shown below:[12]

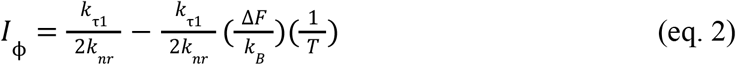

where,

*k*_*nr*_ = *k*_*ic*_ + *k*_*isc*_, *k*_*nr*_, *k*_*ic*_ and *k*_*isc*_ are the rate constants of non-radiative, internal conversion, and intersystem crossing, respectively. Δ*F* refers to the free energy difference between two rotamer states. *k*_τ1_ represents the fluorescence lifetime. Fig.2 presents the energy landscapes and corresponding fluorescence–temperature relationships of a photoactive system with two rotameric states, compared under (a) Δ*F* > 0 and (b) Δ*F* < 0 conditions.

**Figure 2.**
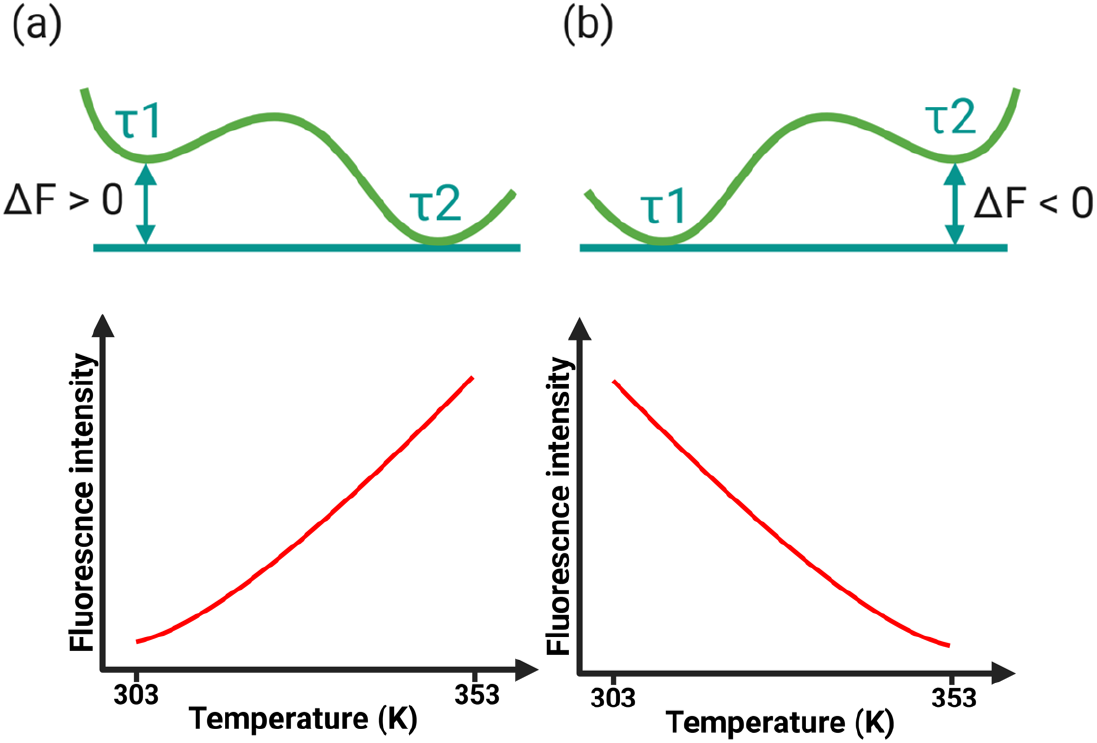
Energy diagrams and the corresponding relationships between fluorescence intensity and temperature for a photoactive/fluorescent system with two rotameric states under (a) F > 0 and (b) F < 0 conditions.

In Fig.2(b), we examine the case where Δ*F* < 0, indicating that the free energy of rotamer state 2 (lifetime τ_2_) is higher than that of rotamer state 1 (lifetime τ_1_). Under this condition, an inverse relationship emerges between fluorescence intensity and temperature. Considering that the Ia^−^ rotamer of Trp134 becomes more populated at elevated temperatures, whereas Ia^+^ persists across all temperatures, we suggest that Ia^+^ corresponds to state 1, while Ia^−^ represents state 2.

### Free energy calculation

Conformational transitions of biomolecules often involve high energy barriers, which hinder efficient sampling within standard molecular dynamics (MD) at room temperature. For instance, in BSA, the intrinsic thermal energy of tryptophan residues may not be sufficient to overcome these barriers, leading to poor sampling of certain rotamers. To enhance sampling along specific conformational coordinates, we employed “umbrella sampling”, which applies a harmonic biasing potential to facilitate exploration across otherwise infrequently sampled regions of phase space.[20] The umbrella sampling simulations were performed using the Collective Variables (Colvars) module implemented in NAMD.[21] The dihedral angle was employed as the collective variable to bias sampling along the rotameric transition coordinate.

The bias potential was defined as:

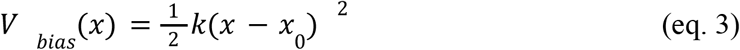

where *x* is the reaction coordinate (a chosen dihedral angle), *x*_0_ is the bias center, and *k* is the force constant. By varying *x*_0_ across a series of windows, the conformational landscape is systematically sampled. In this study, we selected the dihedral angle ∠*C*_α_*C*_β_*C*_γ_*CD*1 (denoted as φ2) to monitor the flipping dynamics of the indole group of Trp134. The angle was sampled from –120° to 120° in 25 windows with 10° spacing, using a harmonic force constant of 0.1 kcal·mol^-1^·Å^-2^. Independent MD simulations were performed for each window to ensure sufficient overlap. (See Fig.S2 of the Supporting Information)

Free energy profiles were then reconstructed using the Weighted Histogram Analysis Method (WHAM)[20]. The resulting potential of mean force (PMF) represents the effective free-energy landscape governing rotameric interconversion along the selected collective variable.[22] WHAM combines the biased histograms from each window and reweights them to recover the unbiased probability distribution, *P*_u_(*x*), from which the potential of mean force (PMF) is obtained:

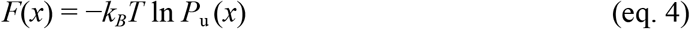

This approach ensures that information from all umbrella windows is optimally combined, reducing statistical error and providing a reliable estimate of the free energy landscape. The optimal estimate of the unbiased distribution is given by,

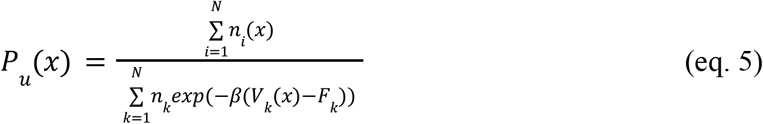

where *n*_i_(*x*) is the number of counts in bin *x* from window *i* (*N* is the number of simulation windows), *n*_*k*_ is the total number of samples from window *k, V*_*k*_(*x*) is the biasing potential applied in window *k*, and *β*=(*k*_*B*_*T*)^−1^. The quantity *F*_k_ is the free-energy offset for window *k*, determined self-consistently to ensure normalization.

### QM/MM Method

QM/MM simulations[23] were performed using the QMMM module implemented in NAMD.[24] System preparation and QM region definition were carried out using the QwikMD interface[25] in VMD (v1.9.4a55).[17]

In this work, we adopted the additive QM/MM scheme.[26] The total energy of the system is expressed as:

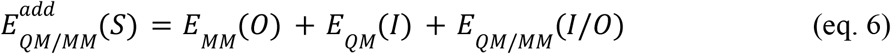

where S, I and O denote the whole system, internal (QM) and external (MM) subsystems, respectively. The QM/MM coupling term is given by:

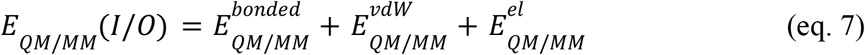

Here,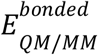 represents bonded interactions between QM and MM boundary atoms, 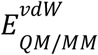 corresponds to van der Waals interactions, and 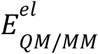 denotes electrostatic interactions.

Unlike the subtractive scheme, in the additive approach, the MM energy is evaluated only for the external subsystem, and an explicit QM–MM coupling term *E*_*QM/MM*_(*I/0*) is included.

For the treatment of the coupling term, we employed electrostatic embedding[26], in which the QM Hamiltonian incorporates the point charges of the external MM region, allowing polarization of the QM electron density by the surrounding environment. See Fig.7(f) for the atomistic details of the QM/MM scheme.

QM calculations were carried out using ORCA (v6.0.0).[27,28] The primary level of theory was B3LYP combined with the def2-SVP basis set,[29] and energies and gradients were evaluated using the ENGRAD scheme. To account for dispersion effects, additional calculations were performed at the B3LYP-D3(BJ)/def2-SVP level of theory. TightSCF convergence criteria were employed, and the RIJCOSX approximation was applied to accelerate exchange integral evaluations. The QM region had a total charge of 0 and a spin multiplicity of 1.

Electrostatic embedding was used for QM/MM coupling, allowing polarization of the QM region by MM point charges. Covalent bonds crossing the QM/MM boundary were treated using the charge-shift (CS) scheme with link atoms (See Fig.7(f)). QM atomic charges were evaluated using Mulliken population analysis.

The MM region was described using the CHARMM36 force fields, which provide bonded and nonbonded parameters for biomolecular systems such as proteins, lipids, and carbohydrates. Water molecules were represented using the TIP3P model. A 12 Å cutoff was applied to nonbonded interactions, with a switching function between 10 and 12 Å. Long-range electrostatics were treated using the particle mesh Ewald (PME) method with a grid spacing of 1 Å.

## RESULTS

### Temperature-induced Rotamer Dihedral Flipping of Tryptophan Reflects Dynamic Response, while Global and Secondary Protein Structures Remain Intact

To evaluate the structural response of BSA to temperature variation, all-atom molecular dynamics (MD) simulations were performed to monitor both global and local conformational changes. The analysis followed a hierarchical framework, examining the backbone structure across three levels: the overall tertiary fold, sub-domain motifs, and individual secondary-structure elements. This multiscale approach provides a systematic view of how temperature affects protein stability and conformational dynamics. Fig.3 shows the root-mean-square deviation (RMSD) profiles of BSA at four different temperatures. At the global level, the average RMSD values remain centered around 4–5 Å, with no significant temperature dependence. Although the RMSD distribution broadens slightly at 353 K, this variation mainly reflects increased thermal fluctuations rather than any irreversible structural deformation.

Focusing on the sub-domain containing Trp134, the mean RMSD decreases to approximately 2 Å with minimal variation across all temperatures (Fig. 3c–d). This finding indicates that the local environment surrounding Trp134 remains structurally stable despite increased thermal motion. Consistent with this observation, the hydrogen-bonding arrangement within the Trp134 subdomain is largely preserved across the crystal structure and MD simulations at both 303 K and 353 K (Fig.S3). Likewise, analysis of individual helices shows negligible RMSD differences among temperatures (Fig. 3e–g), confirming that secondary structures are well preserved. In summary, these results demonstrate that while elevated temperature enhances local dynamical flexibility—particularly relevant to Trp134 rotameric flipping—the overall tertiary and secondary structures of BSA remain intact.

**Figure 3.**
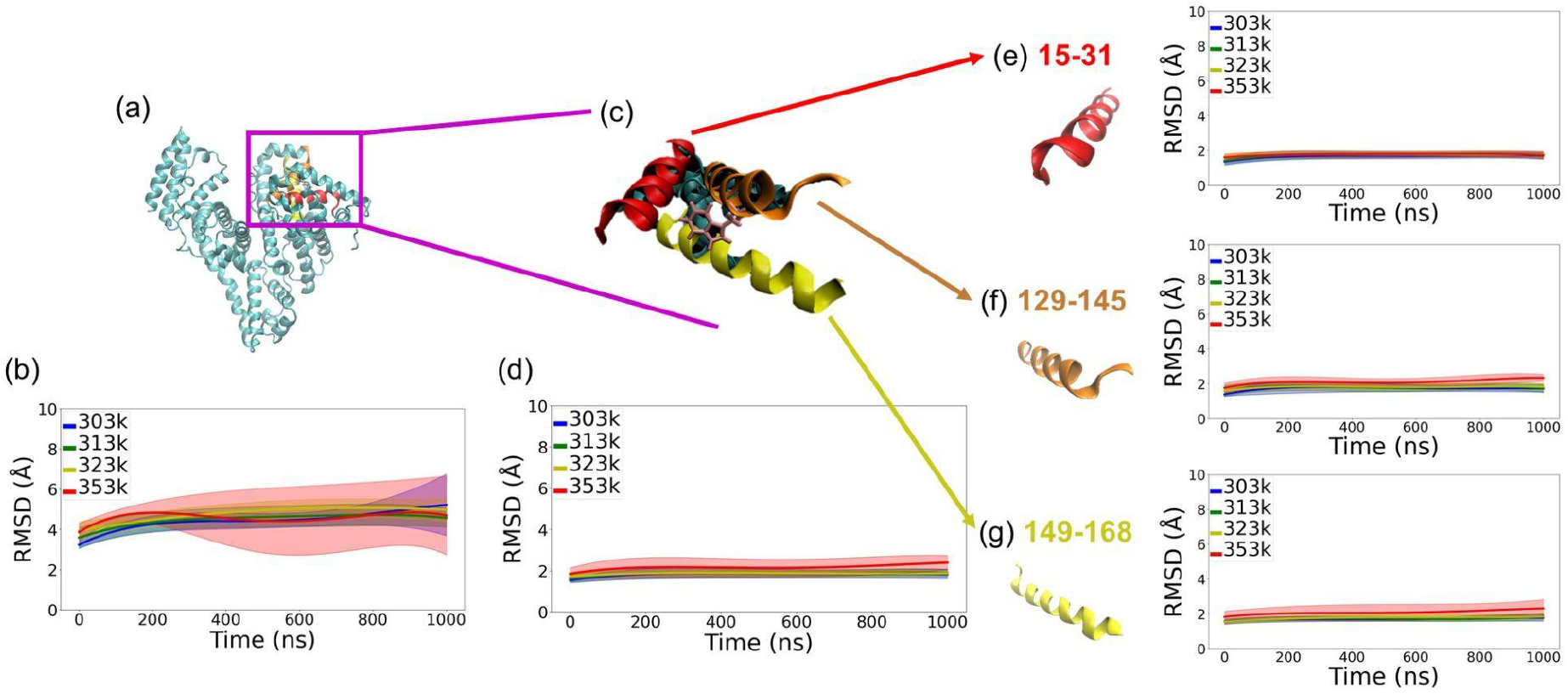
The structural RMSD analysis for the global, sub-domain, and secondary structure of protein BSA at different temperatures is shown. (a) Global structure of (Bovine serum albumin) BSA. Corresponding RMSD analysis as a function of simulation time at different temperatures is depicted in (b). (c) BSA sub-domain consisting of a three-helix bundle (where Trp134 is located in one of the helices), highlighted in the context of the whole BSA protein. The corresponding RMSD analysis is presented in (d). Panels (e-g) represent individual helices (sequence segment: 15-31 a.a. in red, 129-145 a.a. in orange, and 149-168 a.a. yellow, respectively) along with their respective RMSD analyses. The RMSD analyses were carried out for simulations at four different temperatures: 303K, 313K, 323K, and 353K.

Building on this observed structural stability, we next investigate the temperature-dependent dihedral dynamics of Trp134 to elucidate how local rotameric transitions respond to thermal fluctuations. Since the overall protein fold remains intact across all temperatures, the primary temperature-driven effect is expected to arise from internal dihedral flipping rather than global unfolding. The following analysis therefore focuses on quantifying the rotameric transitions of Trp134 and characterizing their associated free-energy landscape to reveal the molecular mechanism underlying temperature-induced fluorescence variation.

### Temperature-Dependent Rotamer Population Redistribution of Trp134

Having established that the overall tertiary and secondary structures of BSA remain stable across the examined temperature range, we next turn our attention to the local conformational dynamics that underlie temperature-dependent fluorescence behavior. In particular, the dihedral motions of Trp134 serve as a sensitive probe for detecting subtle structural responses that do not manifest at the global level. To interpret these local fluctuations quantitatively, we employ a rotamer-based model that links dihedral population shifts to changes in fluorescence intensity. This framework enables us to examine how temperature alters the relative stability and occupancy of Trp134 rotamers, thereby providing a mechanistic explanation for the observed fluorescence variation. Fig.4 summarizes these relationships, highlighting the thermally induced redistribution of rotamer populations and their energetic consequences.

Fig. 4a illustrates the key dihedral pairs—defined by the angles ∠*C*_*COOH*_*C*_α_*C*_β_*C*_γ_and ∠*C*_α_*C*_β_*C*_γ_*CD*1—used to classify Trp134 rotamers (See Methods). The resulting joint distributions of these dihedral angles show how rotamer populations evolve with temperature. By examining population changes among the six possible rotamers (Ia^±^, Ib^±^, Ic^±^), these distributions reveal how thermal fluctuations redistribute population among conformational substates. At lower temperatures, the Ia^+^ is overwhelmingly dominant, whereas increasing temperature progressively shifts the population toward Ia^−^ and minor rotameric states. This temperature-induced Ia^+^ → Ia^−^ transition establishes a direct quantitative link between population dynamics and the observed fluorescence response of tryptophan. Possible contributions from fluorescence resonance energy transfer (FRET) were evaluated and found to be negligible and largely temperature-independent (see Fig.S9 and Fig.S10 Supporting Information), supporting that the observed fluorescence modulation is dominated by rotamer population shifts. The complete temperature-dependent dihedral-angle distributions and corresponding rotamer populations for all simulated temperatures are provided in Fig.S4 of the Supporting Information. To enable a more quantitative comparison, Fig. 4b summarizes the temperature dependence of rotamer populations. As temperature increases, the population of Ia^+^ decreases from nearly 100% to approximately 90%, accompanied by a corresponding increase of the Ia^−^ population from nearly zero to about 10%. The inset highlights this reciprocal trend: a ~10% reduction in Ia^+^ (red curve) is matched by an approximately ~10% increase in Ia^−^ (green curve). In addition, the Ic^+^ and Ic^−^ rotamers (blue and cyan curves) exhibit slight increases with temperature, although their overall populations remain negligible. Fig. 4c shows the population histograms for the Ia^+^, Ia^−^, Ic^+^, and Ic^−^ rotamers at each temperature. Across all conditions, Ia^+^ accounts for more than 90% of the population, dominating the conformational landscape at lower temperatures. At 353 K, however, Ia^−^ emerges as a significant minority population (~10%), accompanied only by trace contributions from Ic^+^ and Ic^−^.

**Figure 4.**
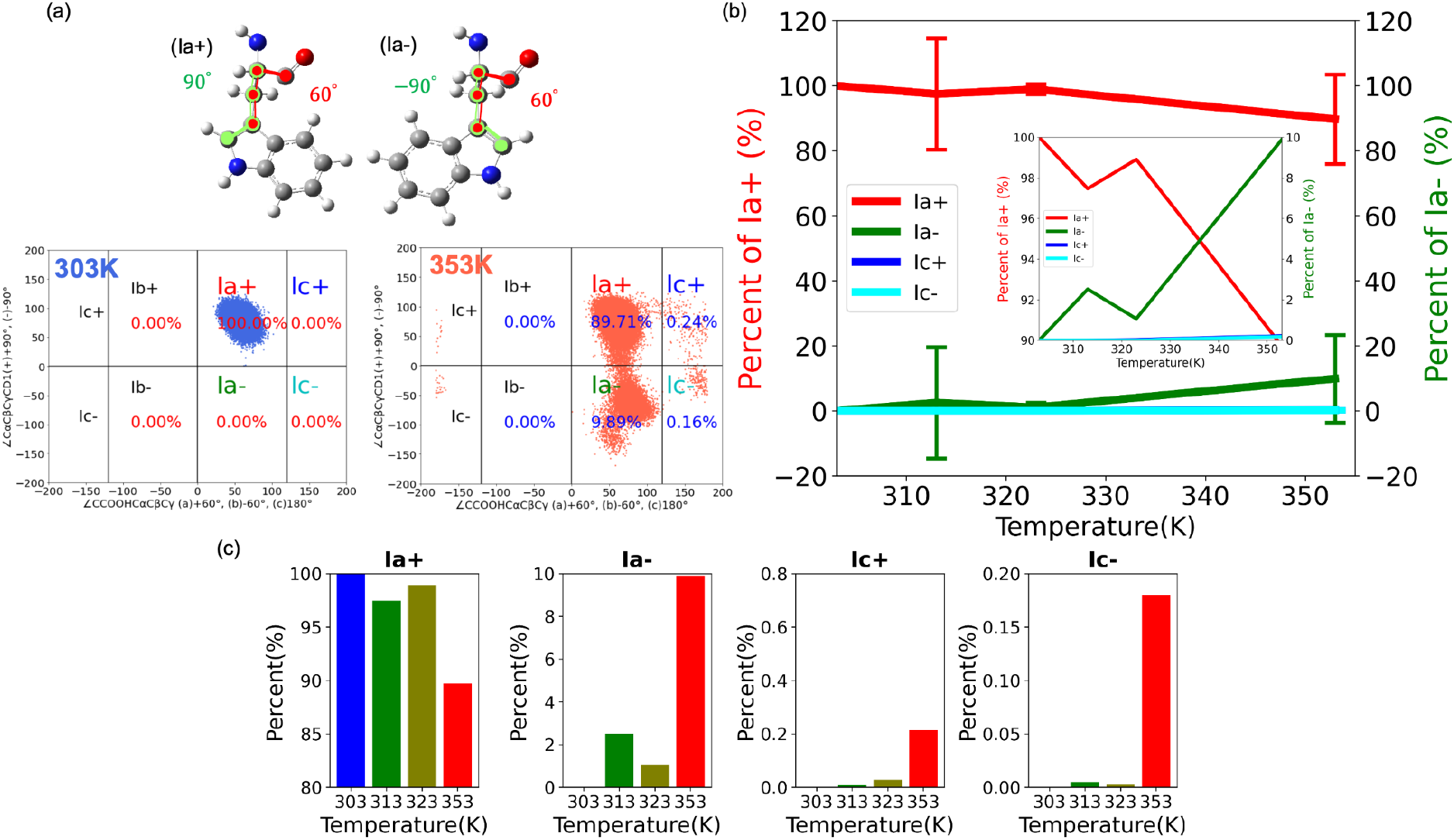
Conformational distributions of tryptophan-134 (Trp134) in BSA at different temperatures. (a) Joint distributions of the dihedral angle pairs ∠*C*_*COOH*_*C*_α_*C*_β_*C*_γ_and ∠*C*_α_*C*_β_*C*_γ_*CD*1 used to classify Trp134 rotamers, shown for 303K (Left) and 353K (Right). (b) Temperature dependence of the population fractions of the four rotamers as a function of temperature. The inset highlights the reciprocal population changes between the Ia^+^ and Ia^−^ rotamers. (c) Population histograms of the Ia^+^, Ia^−^, Ic^+^, and Ic^−^ rotamers (from left to right) at each temperature. Note that different percentage scales are used across the panels.

Taken together, these results demonstrate that temperature-dependent fluorescence modulation arises primarily by reciprocal population shifts between the Ia^+^ and Ia^−^ rotamers. This observation motivates a quantitative analysis of the free-energy landscape governing their relative stability and interconversion.

### Temperature-Dependent Free-Energy Landscape Governing Ia^+^/Ia^−^ Rotamer Stability

To quantify the thermodynamic stability of individual rotamers and characterize their interconversion, we constructed one-dimensional free-energy profiles along the relevant dihedral reaction coordinate. Such profiles provide direct access to relative basin depths and transition-state barriers, thereby offering a thermodynamic interpretation of the observed population shifts. In this study, the ∠*C*_α_*C*_β_*C*_γ_*CD*1 dihedral angle was chosen as the reaction coordinate, and umbrella sampling combined with WHAM analysis (see Methods) was employed to obtain converged potential of mean force (PMF) curves.

Fig.5 compares the free-energy landscapes of Trp134 in BSA at 303 K and 353 K, representing low- and high-temperature conditions, respectively. At both temperatures, the Ia^+^ rotamer corresponds to the global minimum, indicating that it is thermodynamically favored relative to Ia^−^. The relative free-energy difference between Ia^+^ and Ia^−^ is −3.75 ± 0.74 kcal mol^-1^ at 303 K and decreases slightly to −3.33 ± 0.20 kcal mol^-1^ at 353 K. In these profiles, the Ia^−^ basin is defined as the reference (*F* = 0), and the reported values represent the free-energy descent of the Ia^+^ state relative to Ia^−^. Although Ia^+^ remains the dominant state at both temperatures, the reduced Δ*F* at 353 K indicates a relative stabilization of the Ia^+^ basin as temperature increases, consistent with the observed increase in its population.

The activation barrier for the Ia^+^ → Ia^−^ transition also exhibits a modest temperature dependence. At 303 K, the barrier height is 4.47 ± 0.23 kcal mol^-1^, whereas at 353 K it decreases to 4.12 ± 0.11 kcal mol^-1^. This reduction suggests that elevated temperature not only shifts the relative stability of the basins but also lowers the effective barrier separating them, facilitating more frequent interconversion between rotamers.

Beyond changes in barrier height and basin depth, the overall shape of the free-energy landscape evolves with temperature. At 303 K, the PMF exhibits a more sharply defined minimum for Ia^+^ and a comparatively deeper energetic separation from Ia^−^. In contrast, at 353 K the landscape becomes slightly flatter, with a reduced free-energy gap and a smoother profile along the transition region. This flattening reflects increased conformational flexibility at higher temperature and a diminished energetic discrimination between substates. Such landscape reshaping provides a thermodynamic basis for the reciprocal population shifts observed in Fig. 4, linking temperature-dependent fluorescence modulation directly to changes in the underlying free-energy surface.

In short, these results demonstrate that temperature influences both the relative stability of the Ia^+^ and Ia^−^ rotamers and the barrier governing their interconversion. The fluorescence response can therefore be understood as a consequence of subtle yet systematic temperature-induced modifications of the local free-energy landscape.

**Figure 5.**
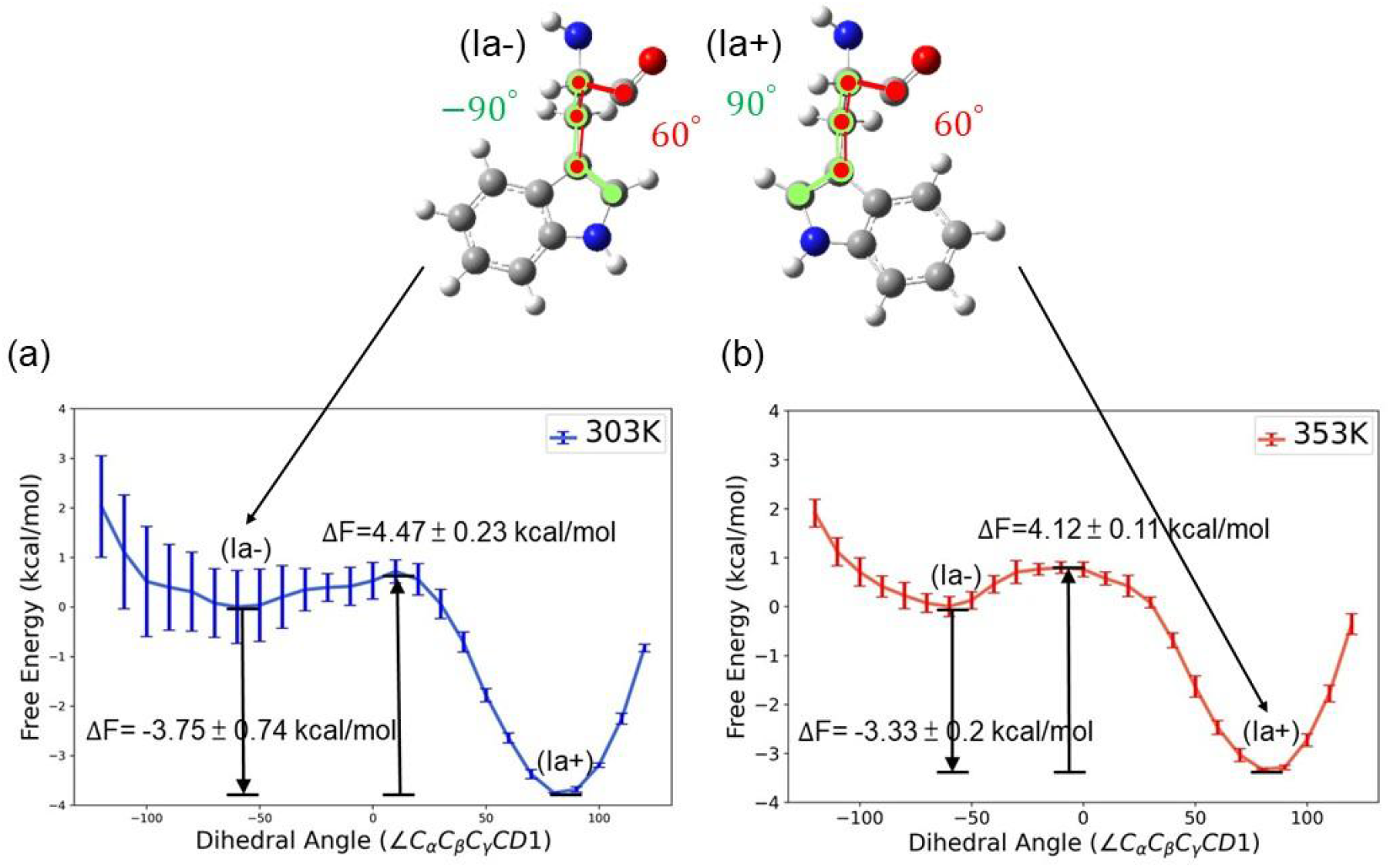
Free energy profile of tryptophan-134 (Trp134) in BSA as a function of the rotamer dihedral angle (∠*C*_α_*C*_β_*C*_γ_*CD*1) at different temperatures. (a) 303 K and (b) 353 K. The potential of mean force (PMF) was obtained from umbrella sampling simulations combined with WHAM analysis. Error bars represent the standard deviation derived from five independent umbrella-sampling datasets. The relative free-energy differences (Δ*F*) are calculated with the Ia^-^ rotamer minimum defined as the reference state (Δ*F* = 0), as indicated in the plots.

### Environmental Modulation of Rotamer Stability and Interconversion Barriers

Having established that temperature reshapes the Ia^+^/Ia^−^ free-energy landscape within BSA, we next examine how different environments modulate the same rotameric transition. By comparing vacuum, aqueous, and protein conditions, we isolate the respective contributions of intrinsic conformational energetics, solvation, and protein confinement to the stability and interconversion of Trp134 rotamers. Fig. 6a and 6b compare the free-energy landscapes of Trp134 in vacuum and aqueous environments. In both cases, the Ia^+^ rotamer remains the most stable state, with the free-energy difference between Ia^+^ and Ia^−^ measured as −1.42 ± 0.50 kcal mol^−1^ in vacuum and −1.51 ± 0.48 kcal mol^−1^ in water. The activation barrier is 2.67 ± 0.48 kcal mol^−1^ in vacuum and decreases to 2.07 ± 0.37 kcal mol^−1^ in aqueous solution, indicating that solvation stabilizes the transition state and effectively lowers the energy barrier. This reduction reflects a general influence of the solvent environment, although determining the specific molecular interactions responsible would require further analysis. We further analyzed hydrogen bonding (see Supporting Information, Fig.S5) and observed a correlation between the average hydrogen bond number and stability, with more hydrogen bonds corresponding to lower free energy, although the differences are small. As a result, the interconversion between rotamers occurs more readily in water, consistent with faster rotameric dynamics under solvated conditions.

**Figure 6.**
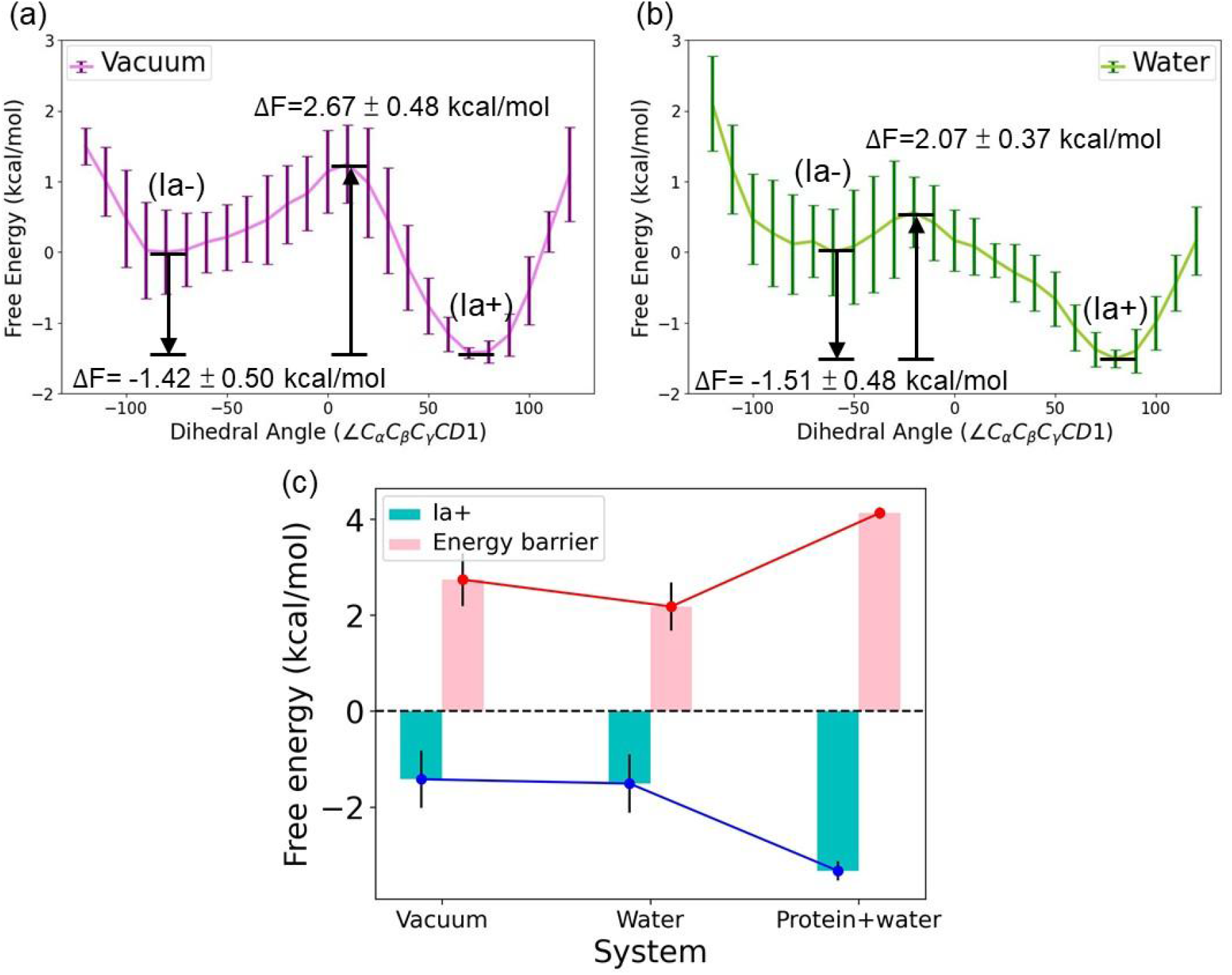
Free-energy landscapes of Trp134 along the ∠*C*_α_*C*_β_*C*_γ_*CD*1 dihedral angle in different environments at 353 K. (a) Gas-phase (vacuum) potential of mean force (PMF). (b) PMF under explicit water solvation. In both panels, the Ia^−^ rotamer minimum is defined as the reference state (*F* = 0), and Δ*F* values denote the relative free energy of the Ia^+^ rotamer. Error bars represent the standard deviation obtained from five independent umbrella-sampling datasets. (c) Summary comparison of the relative free energy of the Ia^−^ state (cyan bars) and the corresponding activation barriers (pink bars) in vacuum, water, and protein (BSA) environments.

Beyond the difference in barrier height, the shapes of the potential of mean force (PMF) profiles also differ markedly between environments. Between the Ia^+^ rotamer and the transition state, the vacuum profile rises sharply, whereas the aqueous PMF increases more gradually, suggesting that solvation smooths the free-energy surface and broadens the transition region, consistent with previous theoretical descriptions of solvent-dependent modulation of biomolecular free-energy landscapes.[30] Similar environment-dependent reshaping of rugged biomolecular free-energy landscapes has been observed in all-atom enhanced-sampling studies, where subtle balances between solvation and local interactions govern conformational stability and transition pathways.[31] Moreover, the transition-state position shifts from a dihedral angle of approximately +10° in vacuum to about −20° in water, implying that solvent molecules influence not only the magnitude but also the location of the barrier along the reaction coordinate.

As more clearly illustrated in Fig.6(c), both the vacuum and aqueous systems exhibit substantially lower activation barriers compared to the BSA environment, highlighting the pronounced structural confinement imposed by the protein matrix. Within BSA, local packing effects, hydrogen bonding, hydrophobic contacts, and other noncovalent interactions collectively stabilize the tertiary structure while restricting the conformational flexibility of Trp134. As a result, rotameric interconversion within the protein requires additional energetic input to overcome these spatial and interaction-based constraints. This observation highlights the important role of the biological environment in modulating the rotamer free-energy landscape and the associated conformational transitions.

### Comparison between MD, QM, and QM/MM calculations

To validate the free-energy differences obtained from molecular dynamics (MD) simulations, we performed quantum-mechanical calculations on an isolated tryptophan residue for comparison. These calculations were carried out both in the gas phase and under implicit solvation using the PCM model. The purpose of this analysis is to provide quantum-level benchmarks against which the MD-derived energetics can be assessed. More broadly, examining whether the population-shift rotamer model of tryptophan is consistent across different levels of electronic structure theory raises an important and fundamental question about the generality of the underlying mechanism.

TD-DFT calculations indicate that the dominant excitation retains localized π–π* character with negligible charge-transfer contribution (Fig.S8). The strong overlap between the hole and particle distributions suggests limited excitation-induced structural relaxation, supporting the use of S0 geometries for qualitative fluorescence analysis. Although the dominant excitation is primarily described by a HOMO→LUMO+1 transition, both orbitals remain localized on the indole aromatic framework, confirming its local-excitation character.

Fig. 7 compares the free-energy profiles calculated using various computational approaches (MD, QM, and QM/MM) under different environmental conditions. For the MD calculations, we separately examine the effects of the protein matrix, explicit water solvation, and vacuum on a single tryptophan residue. For the QM calculations, both gas-phase and dielectric solvation conditions are considered, whereas the QM/MM calculations simultaneously incorporate the protein environment and explicit solvation for benchmarking purposes. The corresponding free-energy differences and activation barriers obtained from all computational approaches are summarized in Table 1. Fig.8 further presents the detailed free-energy profiles associated with each method and environmental condition discussed above.

**Table 1.** Summary of calculated rotamer energetics under different computational and environmental conditions.

| | | $\Delta E^a$<br>(kcal/mol) | $\Delta F$<br>(kcal/mol) | $\Delta S^b$<br>(kcal/K·mol) | Env.<br>(Solvent) | Env.<br>(Protein) |
| --- | --- | --- | --- | --- | --- | --- |
| MD | Protein +<br>solvent | 102.97 | -3.33 | 0.301 | v | v |
|  | Vacuum | -46.499 | -1.42 | -0.128 | x | x |
|  | Solvent | -79.680 | -1.51 | -0.221 | v | x |
| QM | Vacuum | 0.090755 | 0.335 | -0.000692 | x | x |
|  | Solvent<br>(PCM) | 0.180978 | 0.554 | -0.00106 | v | x |

<sup>a</sup>Energy change ( $\Delta E$ ) was calculated using the CHARMM force field as converged sums of mechanical energy. Details are provided in the Supporting Information (See Fig.S6).
<sup>b</sup>Entropy change ( $\Delta S$ ) was calculated using the relation $\Delta S = (\Delta E - \Delta F)/T$ , where $T = 353$ K.

**Figure 7.**
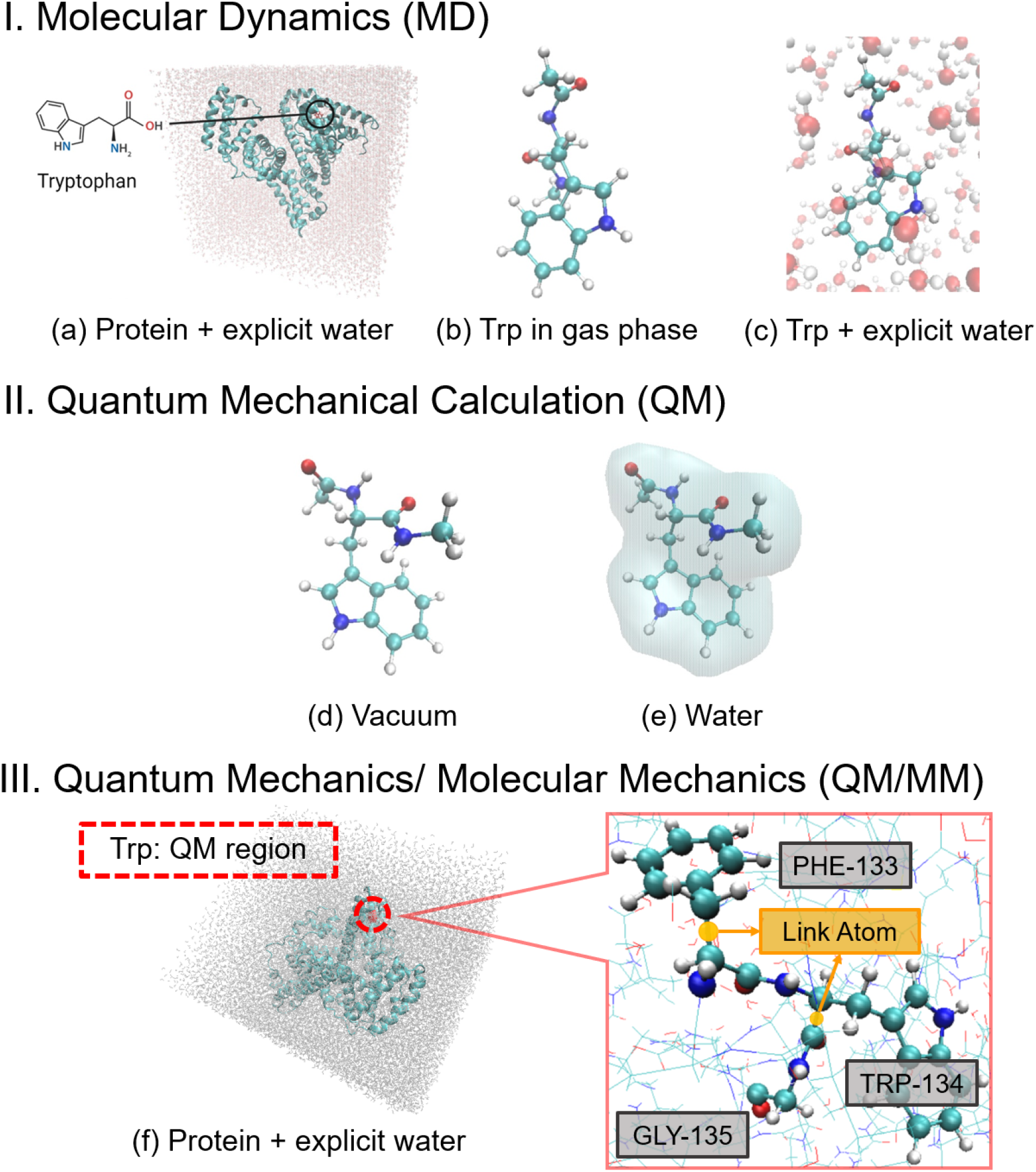
Computational models using molecular dynamics (MD) simulation, quantum (QM) chemistry calculations, and QM/MM at 353K. (a-c) MD. (a) Whole BSA protein in explicit water. (b) Trp in vacuum. (c) Trp in explicit water. (d-e) QM. (d) Trp in vacuum. (e) Solvated Trp (using PCM). (f) QM/MM. Whole BSA protein in explicit water. Two link atoms are introduced: one at the Phe133 backbone between the Cα and carbonyl carbon, and another within the Trp134 backbone between the Cα and carbonyl carbon, where the QM/MM boundary is defined.

**Figure 8.**
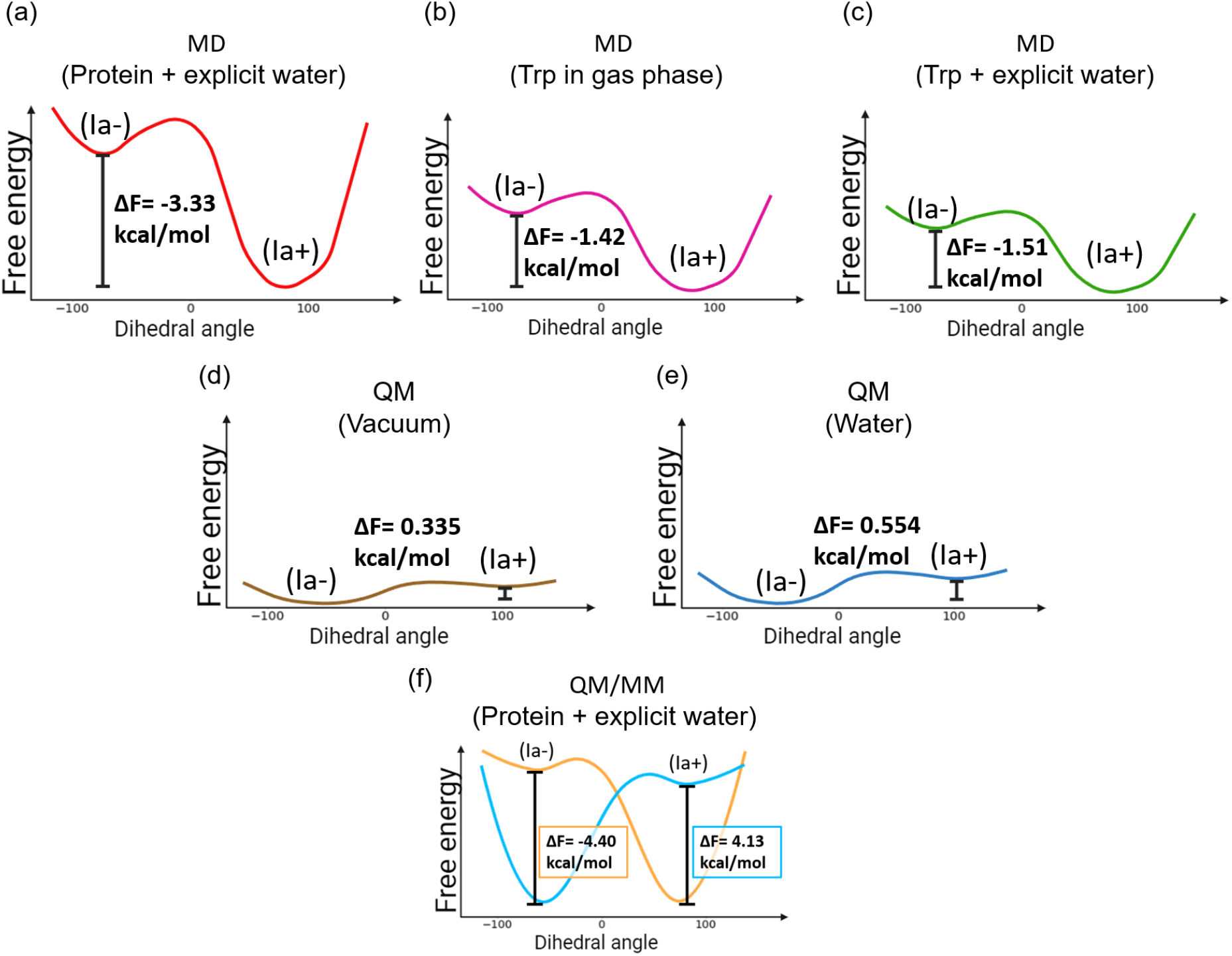
Comparison of the free energy profiles of various computational scenarios using molecular dynamics (MD) simulation, quantum (QM) chemistry calculations, and QM/MM at 353K. (a–f) Correspond to the computational models shown in Fig.7, respectively. The values of Δ*F* are obtained from **Table 1**.

**Figure 9.**
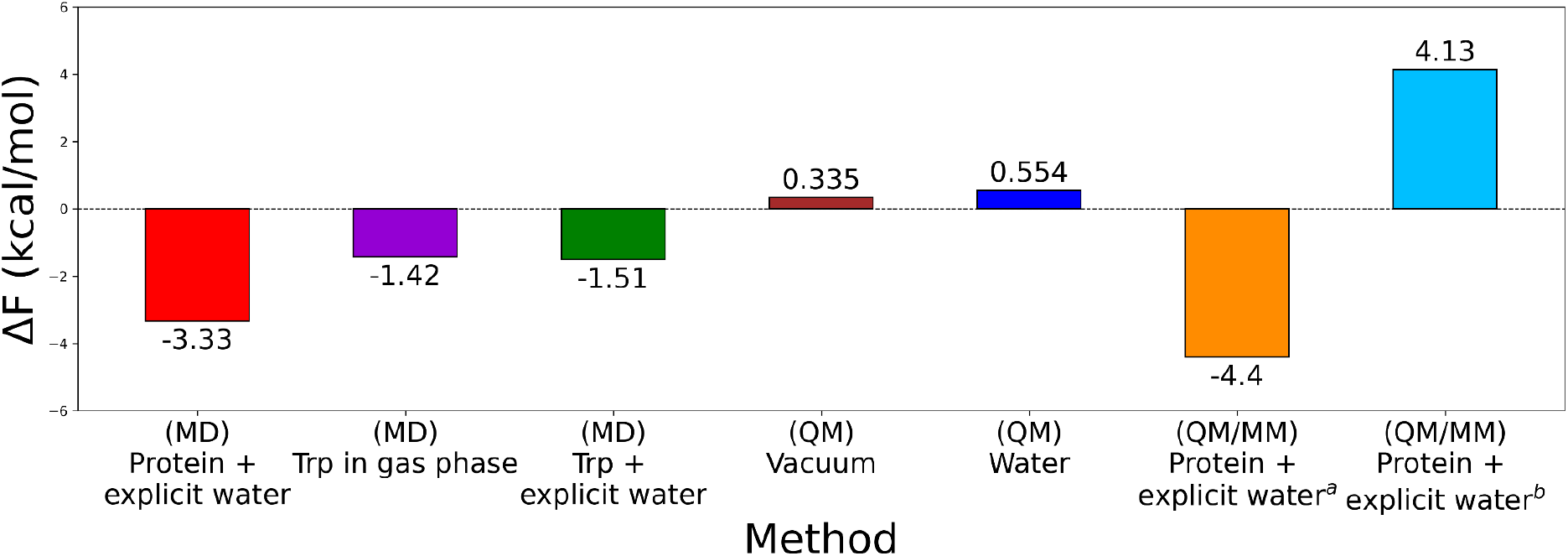
Comparison of free energy differences (ΔF) obtained from molecular dynamics (MD), quantum mechanics (QM), and QM/MM calculations under different environmental conditions. The color scheme used is consistent with that used in Fig.8. In QM/MM, two different basis sets were used for calculation: ^*a*^B3LYP/def2-SVP and ^*b*^B3LYP-D3BJ/def2-SVP with TightSCF and RIJCOSX, both under the protein + explicit water environment.

## DISCUSSIONS

Rotameric population shifts as a general mechanism for temperature-dependent fluorescence

In this study, we propose a generalized molecular mechanism in which the decrease in tryptophan fluorescence quantum yield with increasing temperature is fundamentally governed by population shifts among rotameric conformations of the tryptophan side chain. We suggest that this mechanism is universally applicable across various protein environments, even when other known fluorescence quenching pathways, such as intersystem crossing, photoionization, and intramolecular quenching, coexist.

Previous studies have indicated that with increasing temperature, rotameric populations of tryptophan residues shift due to changes in their relative free energies, leading to a reduction in the proportion of rotamers with higher quantum yields and a corresponding increase in populations with lower quantum yields. This shift consequently manifests as an overall decrease in fluorescence quantum yield.[12] Although additional quenching mechanisms—such as intersystem crossing, photoionization, and intramolecular quenching—may simultaneously occur, their impact typically depends on specific environmental factors, including solvent polarity or pH conditions.

However, experimental evidence shows that even under conditions where intramolecular quenching and other local non-radiative decay pathways are effectively suppressed (e.g., by adjusting pH), the characteristic temperature-dependent decline in tryptophan fluorescence quantum yield remains largely unaffected.[11] Furthermore, the rotameric population shift hypothesis has been theoretically verified under a kinetic framework,[12] complementing the computational results presented in this work. This observation strongly supports the notion that rotameric population shifts constitute a more fundamental and general mechanism. This interpretation is consistent with the broader population-shift framework in biomolecular free-energy landscapes, where subtle energetic perturbations redistribute conformational ensembles and modulate observable functional responses.[13] Thus, we propose that rotameric interconversion acts as the dominant molecular process underlying the temperature dependence of tryptophan fluorescence, operating either independently or synergistically with other quenching pathways. Clarifying this mechanism advances our fundamental understanding of the molecular basis for the temperature sensitivity of tryptophan fluorescence and offers potential implications for future biophysical studies and practical applications.

### Reconciling MD, QM, and QM/MM Descriptions of Rotamer Stability

The comparison between classical molecular dynamics (MD) simulations and quantum mechanical (QM) calculations reveals a clear discrepancy in the predicted trends of tryptophan rotamer stability. MD simulations consistently indicate that temperature-dependent changes in fluorescence intensity arise from population shifts among rotameric states, governed by relatively small free-energy differences that are strongly modulated by the surrounding protein environment. In contrast, standalone QM calculations performed on an isolated tryptophan molecule yield qualitatively different energetic trends, even when implicit solvation effects are included through the PCM model. This discrepancy suggests that the effective free-energy landscape governing rotamer stability is highly sensitive to both environmental constraints and the level of electronic-structure treatment, consistent with previous QM/MM studies emphasizing the role of environmental polarization in biomolecular energetics.[26] Fig.9 summarizes the resulting differences in ΔF obtained from the MD, QM, and QM/MM approaches. The corresponding temperature-dependent quantum-yield trends derived from these ΔF values are provided in Fig.S7 of the Supporting Information.

To reconcile these differences, we employed a QM/MM framework that explicitly accounts for the heterogeneous protein environment while retaining a quantum-mechanical description of the chromophore. QM/MM calculations performed at the B3LYP/def2-SVP level qualitatively reproduce the MD-based picture, correctly capturing the relative stability ordering of key rotamers inferred from temperature-dependent population shifts. This agreement suggests that when the local protein matrix is explicitly included, the rotamer energetics inferred from classical force fields remain physically meaningful at the electronic-structure level. Similar MD–QM/MM approaches have previously demonstrated that biomolecular spectroscopic observables can be directly linked to underlying conformational ensembles and environmental fluctuations.[32]

However, this consistency is not universal across electronic-structure treatments. When dispersion interactions are explicitly incorporated—using B3LYP-D3BJ/def2-SVP with TightSCF and RIJCOSX—the QM/MM results shift toward the trends observed in standalone QM calculations. This sensitivity highlights that the predicted free-energy landscape depends strongly on how noncovalent interactions, particularly dispersion and short-range electronic correlations, are treated. In this context, QM/MM does not simply interpolate between MD and QM descriptions; rather, it reflects a delicate balance between environmental constraints imposed by the protein scaffold and the intrinsic electronic preferences of the chromophore. (See Fig.S7)

These findings emphasize that MD- and QM-based approaches probe different physical aspects of tryptophan fluorescence. Classical MD primarily captures entropic effects, thermal fluctuations, and population redistribution among conformational substates, which are central to understanding temperature-dependent fluorescence behavior. In contrast, QM calculations emphasize electronic energetics and local interaction specificity, which may over-stabilize certain conformations when environmental averaging is insufficiently represented. The divergence between these approaches should therefore not be interpreted as a methodological failure, but rather as a manifestation of the different physical regimes they describe.

Within this framework, rotamer populations emerge as a unifying concept that bridges classical protein dynamics and electronic-structure descriptions of fluorescence. By focusing on population shifts rather than absolute energies, the rotamer model provides a robust, physically interpretable link between temperature-dependent conformational dynamics and observed fluorescence intensity changes. Our results thus underscore the importance of carefully chosen computational frameworks when interpreting fluorescence mechanisms and highlight the value of integrating MD, QM, and QM/MM approaches to achieve a coherent molecular picture.

## CONCLUSIONS

The molecular origin of temperature-dependent tryptophan fluorescence remains an important question in protein photophysics. In this work, we investigated the conformational dynamics of Trp134 in bovine serum albumin (BSA) using molecular dynamics (MD) simulations, free-energy calculations based on umbrella sampling and WHAM, quantum mechanical (QM) calculations, and QM/MM approaches. The MD results show that while the global protein structure remains stable, temperature induces a gradual population shift from the Ia^+^ to the Ia^−^ rotamer, consistent with the experimentally observed decrease in fluorescence intensity. The corresponding free-energy landscapes indicate that this shift originates from subtle changes in basin stability and transition barriers along the rotameric coordinate.

In contrast, standalone QM calculations on isolated tryptophan predict different energetic trends, highlighting the sensitivity of rotamer stability to electronic-structure treatment and the absence of environmental effects. QM/MM calculations partially reconcile these differences by incorporating the protein environment while retaining a quantum description of the chromophore. Together, these results suggest that MD, QM, and QM/MM approaches emphasize complementary physical regimes—thermal population dynamics, intrinsic electronic energetics, and environmental constraints, respectively. In this view, temperature-dependent fluorescence modulation can be interpreted as a population-shift phenomenon occurring on a rugged rotamer free-energy landscape, where subtle energetic perturbations redistribute the equilibrium among metastable conformational substates.[13] Rotamer population therefore emerges as a central physical variable linking protein conformational dynamics with tryptophan photophysics, providing a unified perspective for interpreting temperature-dependent fluorescence in proteins.

## Acknowledgements

This work was supported by the National Science and Technology Council (NSTC), Taiwan, under Grant No. NSTC 113-2628-M-194-001-MY3. The authors acknowledge the National Center for High-performance Computing (NCHC) of the National Applied Research Laboratories (NARLabs), Taiwan, for providing computational and storage resources. The authors are grateful to Prof. Ming-Kang Tsai (NTNU) for insightful discussions on the QM/MM methodology and interpretation of the results. The authors also thank Prof. Yung-Ting Lee (NTTU) for technical support and helpful discussions regarding the computational assessment of fluorescence resonance energy transfer (FRET) effects.

## Graphical Abstract Caption (for CPL)

“Temperature reshapes the rotamer free-energy landscape of Trp134, leading to population shifts that modulate intrinsic tryptophan fluorescence in proteins.”

## Declaration of generative AI and AI-assisted technologies in the manuscript preparation process

During the preparation of this work the author(s) used ChatGPT in order to improve the writing of the manuscript. After using this tool/service, the author(s) reviewed and edited the content as needed and take(s) full responsibility for the content of the published article.

## Notes

### Competing Interest Statement

The authors have declared no competing interest.

### Summary of Updates

References updated; Supplemental files updated; author list updated; acknowledgement updated

