## Supporting Info for "Temperature-Dependent Rotamer Population Shifts Govern Tryptophan Fluorescence in Proteins"

**Supporting Information** 1

Temperature-Dependent Rotamer Population Shifts Govern Tryptophan Fluorescence in Proteins 1

Figure S1. Structural models and RMSD analyses of BSA under different ionic concentrations. 2

Figure S2. Example distribution of umbrella sampling trajectories. 3

Figure S3. Hydrogen-bonding arrangements within the Trp-134 subdomain at different structural conditions. 3

Figure S4. Temperature-dependent dihedral-angle distributions and rotamer populations of Trp-134 obtained from MD trajectories. 4

Figure S5. Free-energy landscapes of Trp134 along the ∠CCCCD1 dihedral angle in water at 353 K and hydrogen analysis. 5

Energy Change (ΔE) Calculation in Table 1 6

Figure S7. The relationship between the quantum yield and the temperature of Trp134 in different systems (MD, QM and QM/MM). 7

Figure S8. Molecular orbital (MO) distributions and corresponding orbital energies of tryptophan calculated at the $S_0$ and $S_1$ geometries under different environments: (a) vacuum and (b) PCM implicit solvent model. 8

Evaluation of FRET Contributions to Tryptophan Fluorescencer 9
Figure S9. The structural distribution of Trp and Tyr within BSA. 11
Figure S10. Fluorescence intensity distributions of TRP–TYR pairs in BSA as a function of time at 303 K and 353 K. 12

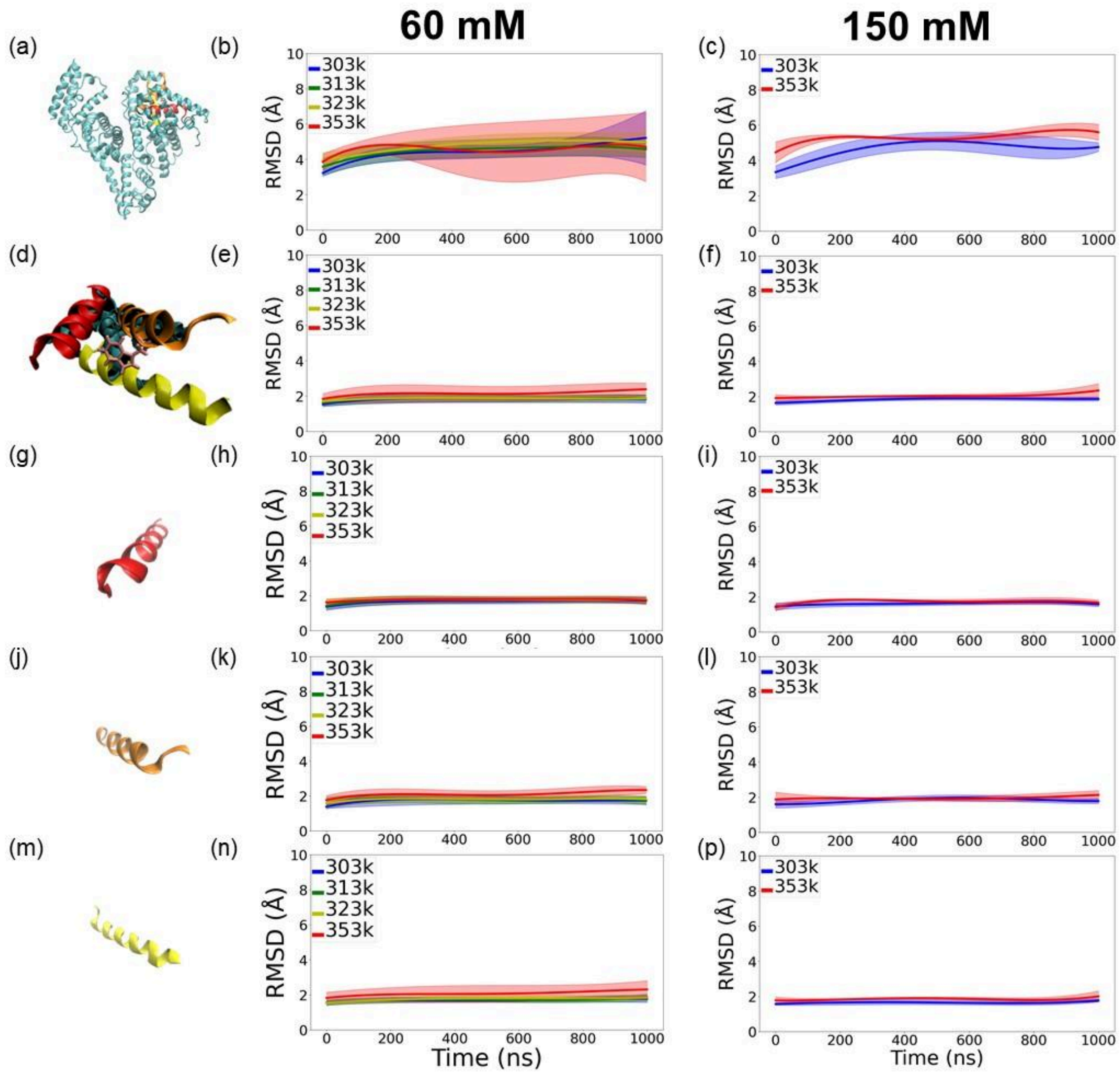


**Figure S1. Structural models and RMSD analyses of BSA under different ionic concentrations.**

The protein was solvated in a water box and neutralized with 17 $Na^+$ ions, corresponding to an ion concentration of 60 mM. Additional simulations were performed at 150 mM to mimic physiological ionic strength conditions. The structures shown here are identical to those presented in the main text. Panels (a,d,g,j,m) display the structural regions analyzed, including the global structure of BSA, the Trp-134-containing three-helix bundle subdomain, and the individual helices. Panels (b,e,h,k,n) and (c,f,i,l,p) present the corresponding RMSD analyses at 60 mM and 150 mM, respectively. The three individual helices correspond to residues 15–31 (red), 129–145 (orange; containing Trp-134), and 149–168 (yellow). The RMSD profiles indicate that the local helical structures remain relatively stable throughout the simulations under both ionic strength conditions.

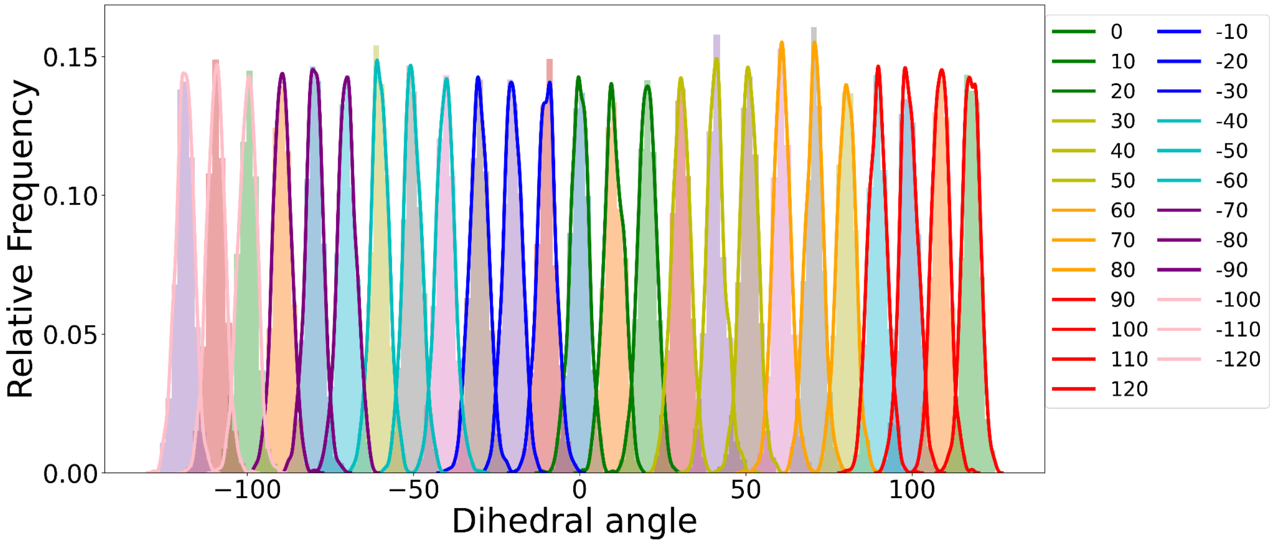


**Figure S2. Example distribution of umbrella sampling trajectories.**

Each curve represents an independent umbrella sampling window, arranged from left to right according to the bias center positions ranging from –120° to 120°, resulting in a total of 25 windows. The labels in the legend denote the bias centers ($x_0$) used in the harmonic biasing potential defined in eq. 3 of the main text. The overlap between neighboring windows ensures sufficient configurational continuity for subsequent WHAM free energy reconstruction.

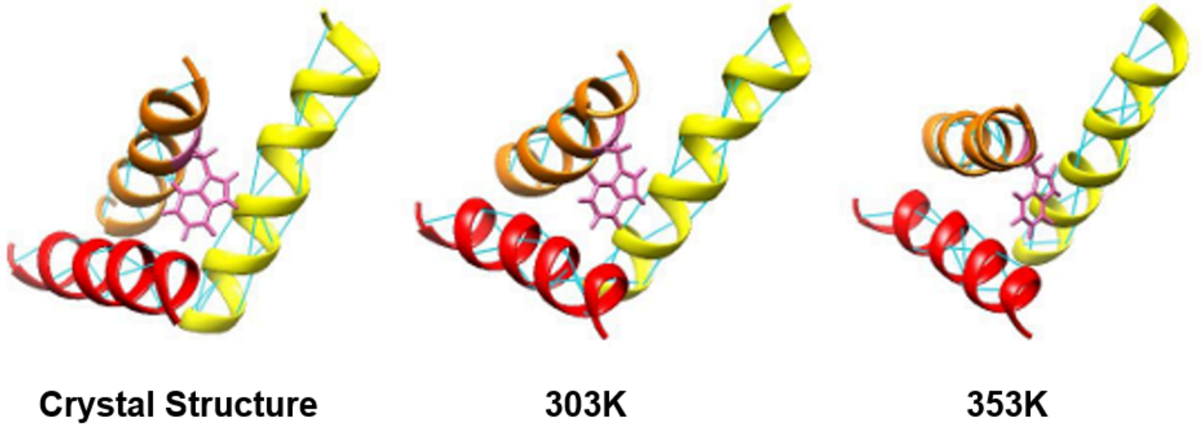


**Figure S3. Hydrogen-bonding arrangements within the Trp-134 subdomain at different structural conditions.**

The hydrogen-bonding networks (blue dashed lines) are shown for the (left) crystal structure, (middle) MD simulation at 303 K, and (right) MD simulation at 353 K. The helices corresponding to residues 15–31, 129–145, and 149–168 are colored red, orange, and yellow, respectively. Trp-134 is highlighted in a magenta licorice representation.

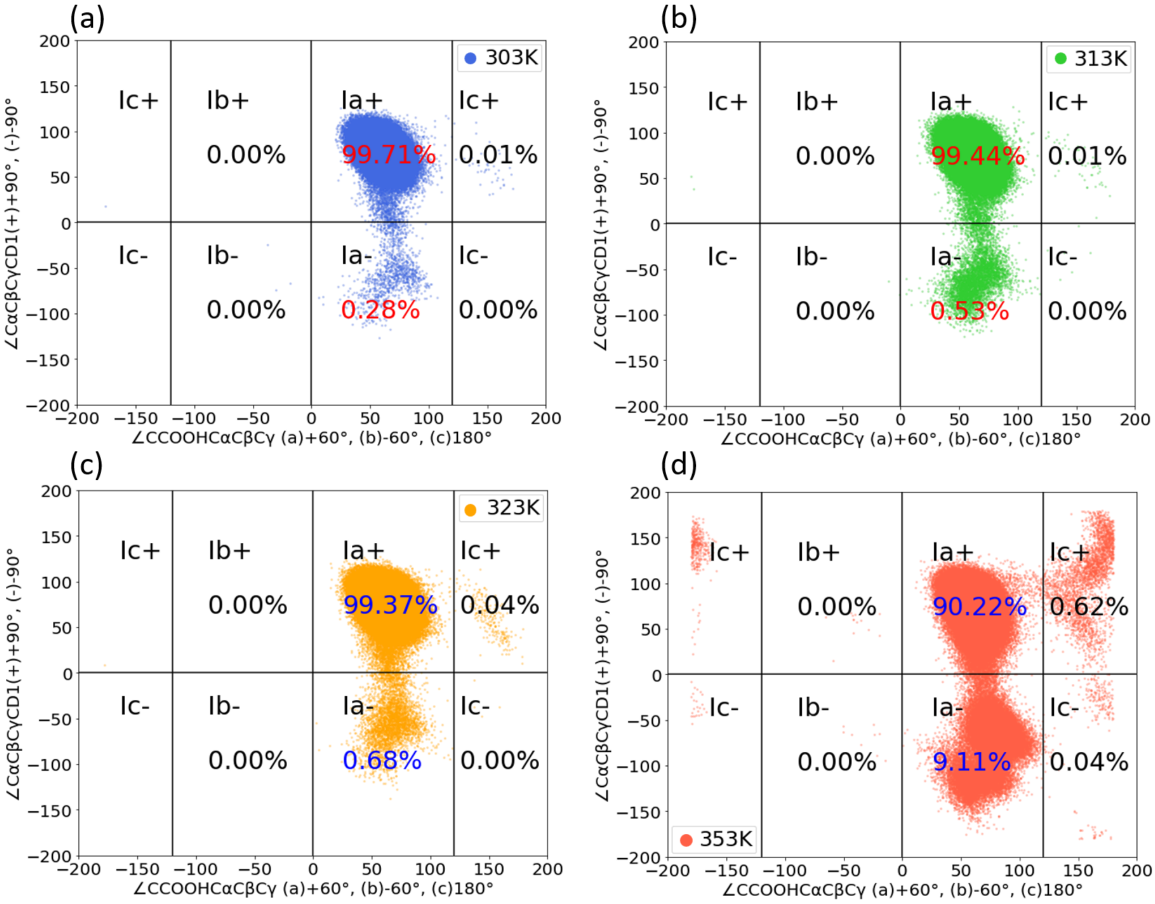


**Figure S4. Temperature-dependent dihedral-angle distributions and rotamer populations of Trp-134 obtained from MD trajectories.**

The joint dihedral-angle distributions and corresponding rotamer population percentages are shown for simulations performed at (a) 303 K, (b) 313 K, (c) 323 K, and (d) 353 K. The results illustrate the progressive redistribution of rotamer populations with increasing temperature.

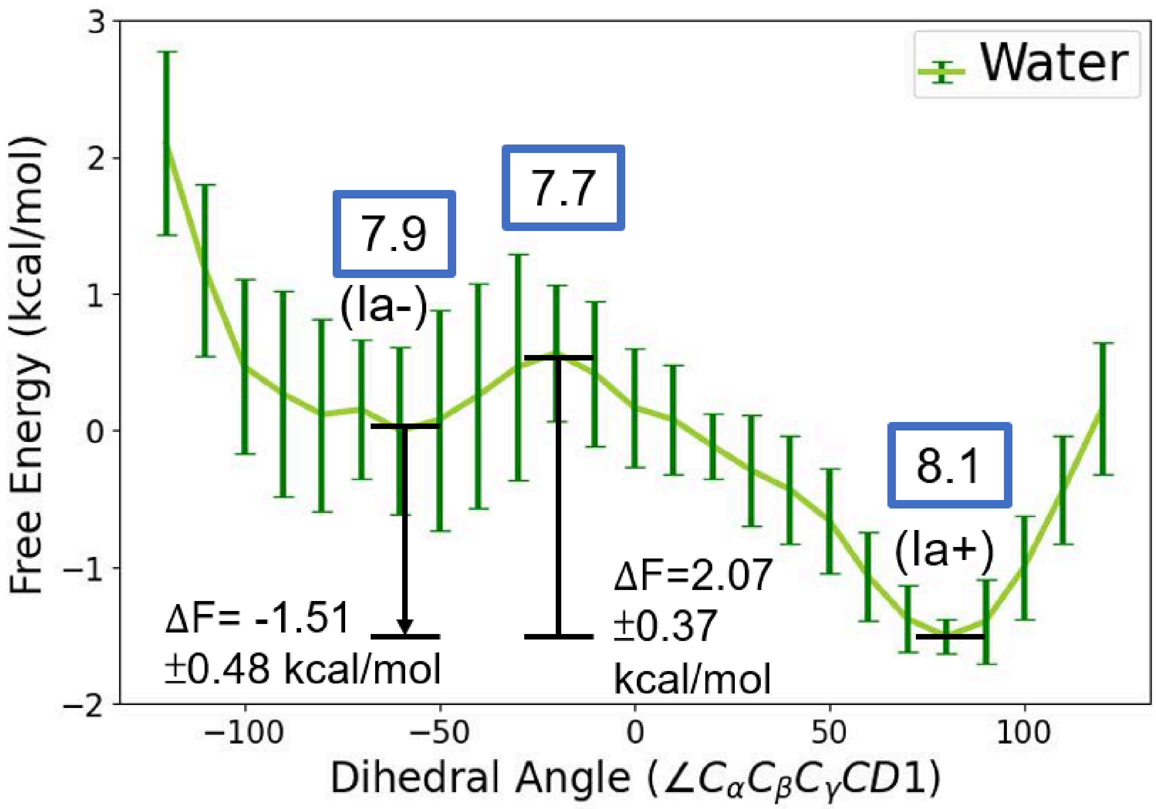


**Figure S5. Free-energy landscapes of Trp134 along the $\angle C_\alpha C_\beta C_\gamma CD1$ dihedral angle in water at 353 K and hydrogen analysis.**

Same as Figure 6(b), showing the free energy profile of the dihedral angle $\angle C_\alpha C_\beta C_\gamma CD1$ of Trp-134 in water. The values highlighted in blue boxes denote the average number of hydrogen bonds formed between Trp-134 and surrounding water molecules within each dihedral region. A general correlation between hydrogen bonding and stability is observed, where conformations with a higher number of hydrogen bonds tend to exhibit lower free energy, although the differences in hydrogen bond counts are relatively small.

Hydrogen bonds were calculated using the hydrogen bond analysis implemented in VMD. For each characteristic region along the dihedral coordinate ($Ia^+$, $Ia^-$, and the transition state), three representative angles were selected, consisting of the central angle and two neighboring points. For example, the transition state is located at approximately 20°, and the hydrogen bond count was obtained by averaging the values calculated at 10°, 20°, and 30°. The same procedure was applied to the $Ia^+$ and $Ia^-$ regions to ensure consistent statistical sampling.

#### Energy Change (ΔE) Calculation in Table 1

To enable a direct comparison with Trp-134, energies (ΔE) were evaluated using the NAMD Energy plugin in VMD. In the whole BSA protein in the explicit water system, spherical regions centered on the center of mass of Trp-134 were defined with radii of 1, 2, 3, 4, 5, 6, 8, 10, 12, 14, and 16 Å. At each cutoff distance, the interaction energy between Trp-134 and the surrounding atoms within the defined sphere was calculated, shown in Fig. S5 below.

We observed that the interaction energy converges when the cutoff distance ranges from 5 to 12 Å. Therefore, ΔE was determined by averaging the energies obtained within this convergence region.

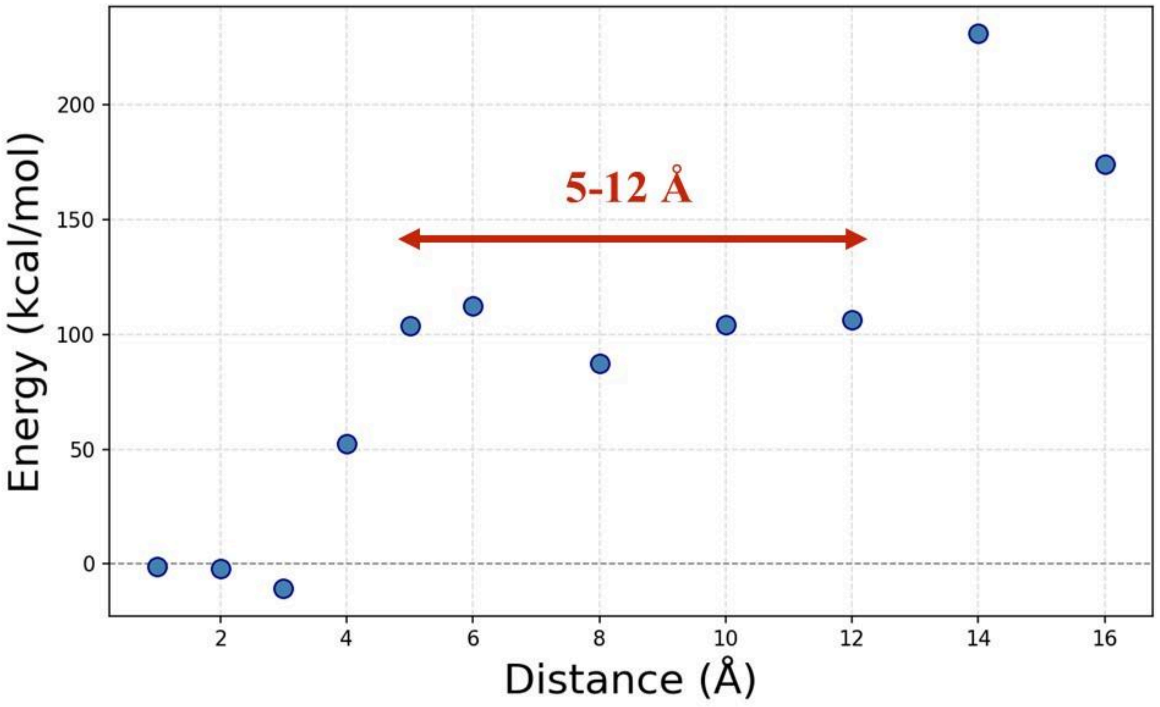


**Figure S6. Energy change (ΔE) between Trp-134 and surrounding atoms as a function of cutoff distance in BSA. The energy shows convergence within the 5–12 Å range.**

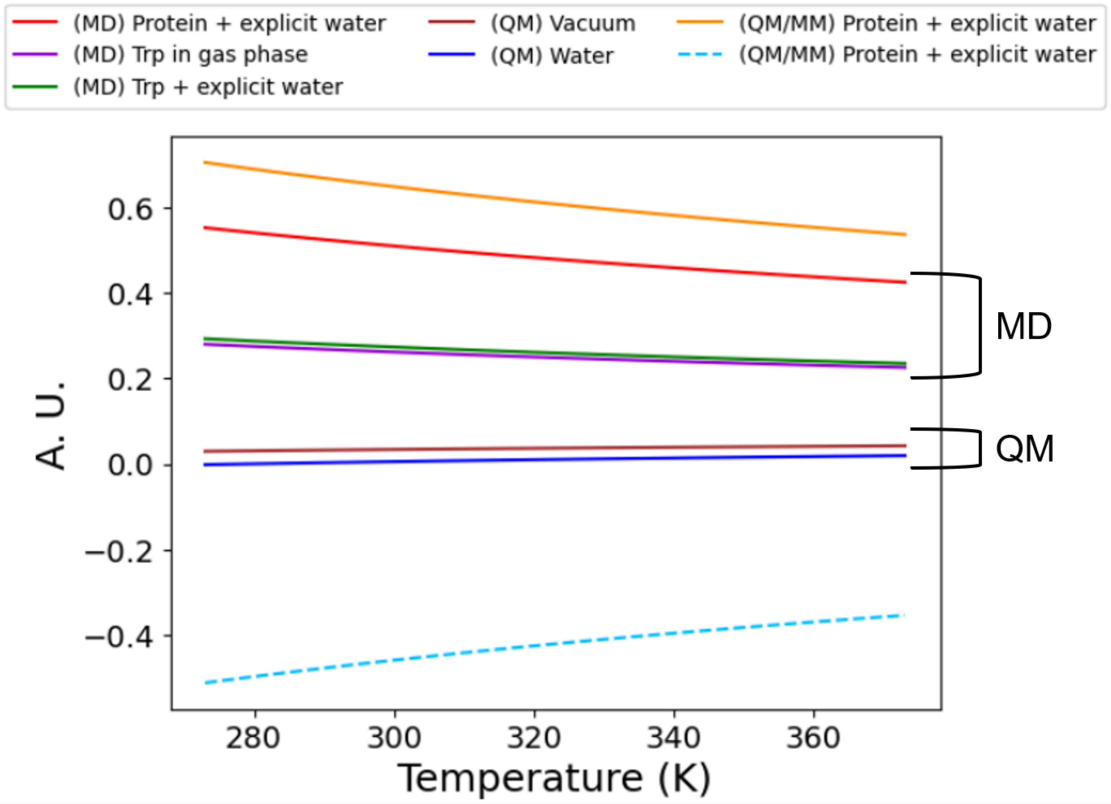


**Figure S7. The relationship between the quantum yield and the temperature of Trp134 in different systems (MD, QM and QM/MM).**

Comparison of quantum yield trends derived from MD, QM, and QM/MM calculations using Eq. (2), highlighting the distinct roles of environmental effects and electronic-structure treatments in determining tryptophan rotamer energetics (quantum yield is shown in an arbitrary unit, A.U.). In all cases, the sign and magnitude of $\Delta F$ serve as the primary determinants of the temperature dependence of quantum yield (solid lines). See Fig. 10 for the value of $\Delta F$ in the histogram. A representative QM/MM-derived profile is additionally shown as a dashed line for comparison; the corresponding quantum yield values are not intended to be quantitatively accurate, but are included to illustrate the qualitative impact of the sign of $\Delta F$ on the predicted temperature-dependent behavior.

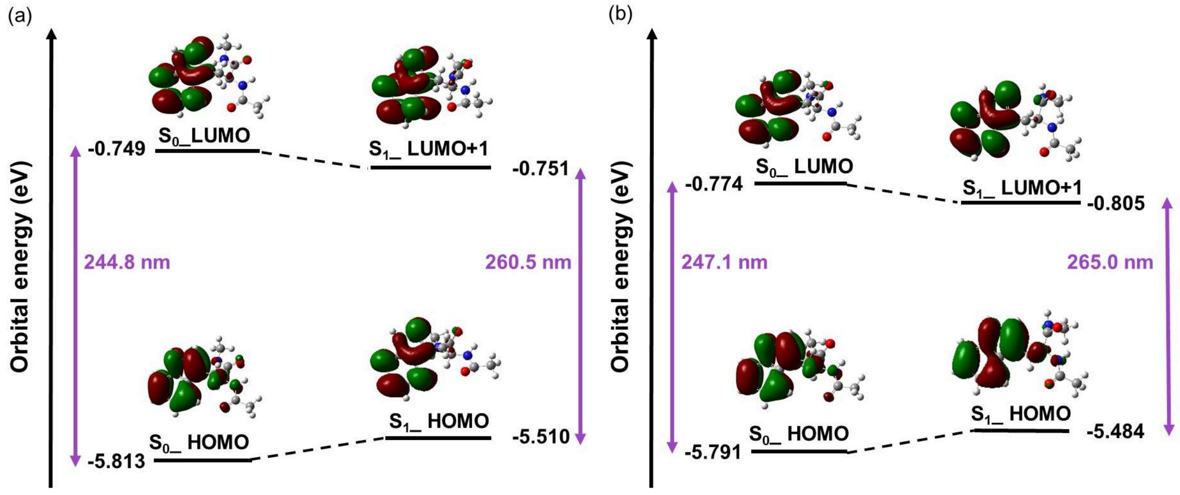


**Figure S8. Molecular orbital (MO) distributions and corresponding orbital energies of tryptophan calculated at the $S_0$ and $S_1$ geometries under different environments: (a) vacuum and (b) PCM implicit solvent model.**

The dominant excitation is primarily described by a HOMO→LUMO+1 transition, with both orbitals localized on the indole aromatic framework. The occupied and virtual orbitals exhibit clear π and π* character, respectively, as evidenced by their delocalized distributions and the increased nodal structure of the virtual orbital. Comparison of the MO patterns and energy levels at the $S_0$ and $S_1$ geometries reveals only minor changes upon excitation, indicating that the electronic structure of tryptophan is only weakly perturbed in the excited state. Together with the strong spatial overlap between the involved orbitals, these results support a localized π–π* excitation with negligible charge-transfer character and justify the use of ground-state geometries for qualitative analyses of fluorescence-related electronic properties.

### Evaluation of FRET Contributions to Tryptophan Fluorescencer

*Fluorescence Resonance Energy Transfer (FRET)* is a non-radiative process in which electronic excitation is transferred from a donor fluorophore to an acceptor fluorophore in a distance-dependent manner, without the emission of a photon. The efficiency of FRET decreases with the sixth power of the donor–acceptor separation, making it highly sensitive to intermolecular distances in the range of 10–100 Å.

In bovine serum albumin (BSA), intrinsic tryptophan (Trp) residues serve as natural fluorescence donors. Neighboring aromatic residues, such as tyrosine (Tyr) and phenylalanine (Phe), can act as potential acceptors. When these residues are spatially close to Trp, FRET may occur, resulting in quenching of Trp fluorescence intensity and modification of its fluorescence lifetime. Such effects can complicate the interpretation of experimental data. Therefore, when analyzing Trp fluorescence in BSA, it is essential to consider possible FRET contributions in order to disentangle intrinsic rotameric effects from intermolecular energy transfer.

The efficiency of energy transfer, *E*, is given by:

$$E = \frac{1}{1+\left(R/R_0\right)^6} \qquad \text{(S1)}$$

where $R$ is the donor–acceptor separation and $R_0$ is the Förster radius, defined as the distance at which $E$=0.5. The donor intensity decay function is defined as

$$I_{DA}(t) = \int_0^\infty P(r)I_{DA}(r,t)dr = I_{DA}^0 \int_0^\infty P(r)exp\left[-\frac{t}{\tau_D} - \frac{t}{\tau_D}\left(\frac{R_0}{r}\right)^6\right]dr \qquad \text{(S2)}$$

This is the equation for the **total fluorescence intensity decay of the donor over time**.

In Eq.(S2), $I_{DA}(t)$: The total fluorescence intensity of the donor at time *t*; $I_{DA}(r, t)$: The fluorescence intensity decay at time *t* for a donor-acceptor at distance *r*; $I^0_{DA}$: The initial fluorescence intensity of the donor (at time *t*=0); $P(r)$: The probability distribution function for the donor-acceptor at distance *r*, representing the distribution of distances between the two molecules; $\tau_D$: The donor lifetime, which is the natural decay time without energy transfer. $R_0$: The Förster distance, at which the energy transfer efficiency between the donor and acceptor is 50%; $r$: The actual distance between the donor and acceptor molecules. Note that the fluorescence intensity decay at time *t* for a donor-acceptor at distance *r* is given by

$$I_{DA}(r, t) = I^0_{DA} exp\left[-\frac{t}{\tau_D} - \frac{t}{\tau_D}\left(\frac{R_0}{r}\right)^6\right], \quad \text{(S3)}$$

and the **Gaussian distribution function** *P*(*r*) is represented by

$$P(r) = \frac{1}{\sigma\sqrt{2\pi}} exp\left[-\frac{1}{2}\left(\frac{\bar{r}-r}{\sigma}\right)^2\right]. \quad \text{(S4)}$$

It represents the probability of finding the donor and acceptor molecules at a particular distance.

Given the FRET pairs formed between Trp and Tyr residues within BSA, we assess their mutual distance separation, as shown in Fig. S9. Fig. S10 presents the simulated fluorescence decay as a function of time for Trp-Tyr at different temperatures.

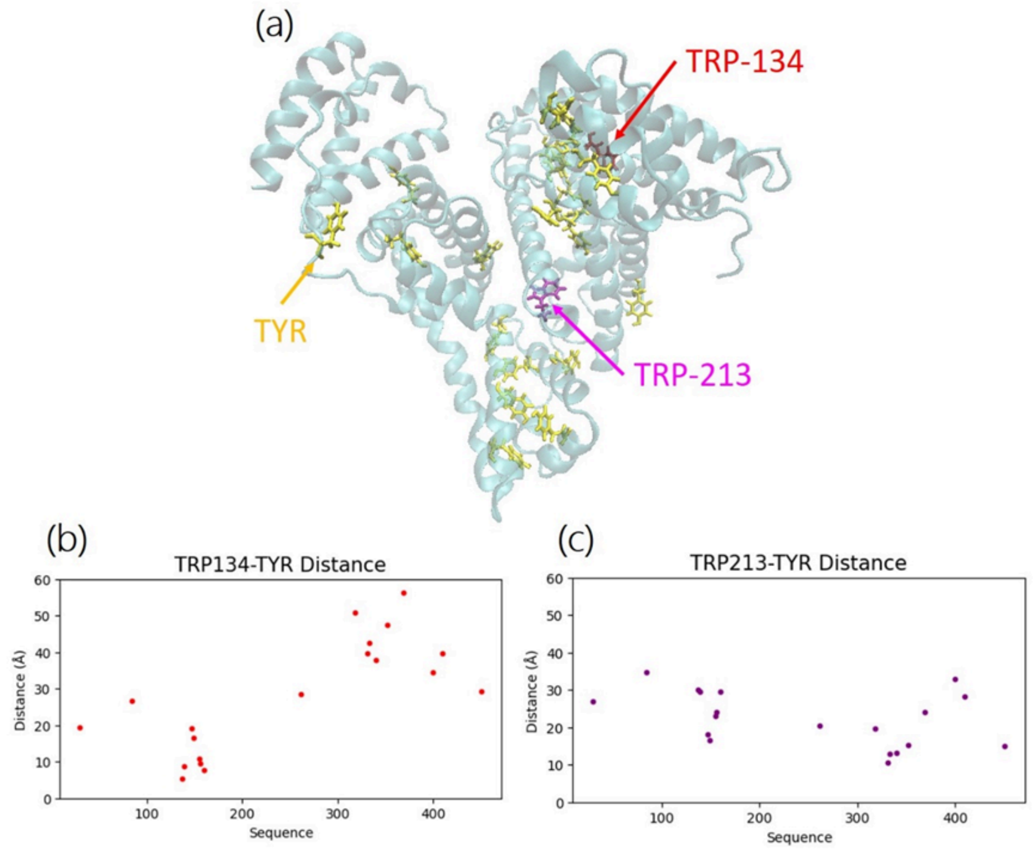


**Figure** S9. The structural distribution of Trp and Tyr within BSA.

(a) Structure of tryptophan (Trp) and tyrosine (Tyr) residues in BSA, and distance distributions between Tyr (yellow) and (b) Trp-134 (red) and (c) Trp-213 (pink). Note that Trp-Phe is not considered due to its low FRET efficiency.

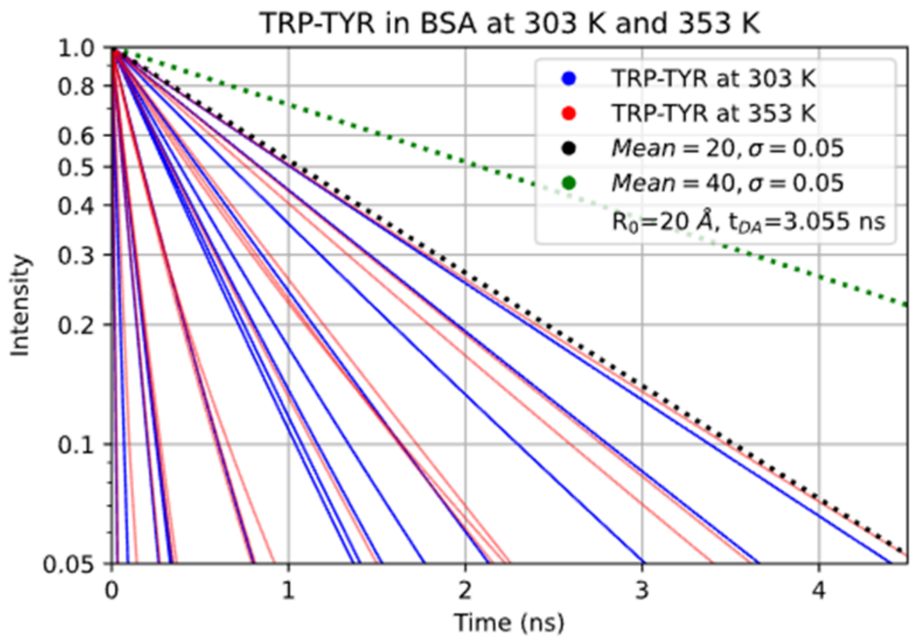


Figure S10. Fluorescence intensity distributions of TRP–TYR pairs in BSA as a function of time at 303 K and 353 K.
